# ExplainBind: Explainable Physicochemical Determinants of Protein–Ligand Binding via Non-Covalent Interactions

**DOI:** 10.64898/2026.03.03.707476

**Authors:** Zhaohan Meng, Zhen Bai, Ke Yuan, Jiangning Song, Yumeng Zhang, Jaime H. Cheah, Wei Jiang, Adam Skepner, Kevin J. Leahy, Iadh Ounis, William M. Oldham, Zaiqiao Meng, Hao Xu, Joseph Loscalzo

## Abstract

Protein-ligand binding governs enzymatic catalysis, metabolic homeostasis, and therapeutic modulation. Thus, the accurate prediction of these interactions underpins modern rational drug discovery. However, existing deep-learning frameworks largely operate as black-box predictors that fail to resolve the individual residues mediating binding or decode the fundamental non-covalent forces that drive molecular recognition. To address these limitations, we present **ExplainBind**, an interaction-aware framework that predicts binding likelihood, localizes specific binding residues at single-amino-acid resolution rather than coarse pocket-level regions, and decodes the underlying non-covalent interaction patterns, all zero-shot, without requiring prior three-dimensional structural inputs. To support residue- and interaction-level training and evaluation, we construct **InteractBind**, a protein–ligand benchmark with residue–atom interaction maps. Simultaneously, benchmarking experiments demonstrate that ExplainBind consistently outperforms state-of-the-art baselines across diverse protein and ligand spaces, maintaining high precision when generalized to entirely novel sequences and chemical scaffolds. When applied to two unseen therapeutic targets, ExplainBind successfully ranks potent angiotensin-converting enzyme (ACE) inhibitors and clarifies differences in their potency via affinity-stratified interaction landscapes. Furthermore, we demonstrate the prospective utility of ExplainBind by discovering novel inhibitors and activators of L-2-hydroxyglutarate dehydrogenase (L2HGDH) through wet-lab validation, with mechanistically distinct interaction profiles providing a clear molecular rationale for their divergent functional outcomes. Collectively, these results establish ExplainBind as a powerful, generalizable tool for mechanistically informed, interpretable drug discovery.

## 1 Introduction

The machinery of life is orchestrated through intricate networks of molecular interactions [1, 2]. Enzymes accelerate chemical reactions by stabilizing transition states, signaling proteins transmit information through transient contacts, and therapeutics achieve efficacy by engaging precise binding sites [3]. Underlying these processes is a constellation of non-covalent forces: hydrogen bonds, salt bridges, van der Waals contacts, hydrophobic contacts, *π*–*π* stacking, and cation–*π* interactions that collectively dictate affinity, specificity, and kinetics [4]. Acting in concert, these interactions stabilize enzymatic catalysis, organize macromolecular assemblies, and confer selectivity to drugs [5]. Crucially, they are exquisitely sensitive to select structural determinants: the loss of a single hydrogen bond, the reorientation of an aromatic ring, or the disruption of a hydrophobic patch can shift a system from active to inert, or from selective to promiscuous. Non-covalent interactions are, therefore, not incidental features, but the fundamental determinants of molecular recognition across biology and medicine [6].

Capturing this physical precision has long demanded three-dimensional structures, where atomic geometries directly encode the non-covalent forces governing molecular recognition. Classical physics-based approaches, molecular docking [7] and molecular dynamics simulations [8], model interaction energies directly from atomic coordinates to rationalize binding and specificity. More recently, geometric deep learning has extended this paradigm to data-driven settings: MaSIF [9, 10] learns molecular surface fingerprints at protein–protein interfaces, and related ideas have since given rise to distance-aware geometric models for protein–ligand docking [11–14]. Yet, in these approaches, distance-based geometry serves only as an implicit proxy for molecular interactions, the underlying non-covalent interactions are never directly represented. This limitation runs deeper than it first appears: recent adversarial analyses of representative protein–ligand co-folding models, including AF3 [15], RoseTTAFold All-Atom [16], Chai-1 [17] and Boltz-1 [18], reveal that high structural accuracy does not guarantee physically meaningful interactions — systematic violations of steric and electrostatic principles persist even at low root-mean-square deviation (RMSD), exposing a fundamental discordance between geometric fidelity and chemical realism [19].

This gap is not confined to structure-based models but extends equally to sequence-based approaches [20–24], where the field has converged on protein–ligand binding (PLB) prediction as a scalable surrogate objective [25, 26]. Sequence-based PLB models typically employ cross-attention or related fusion modules and claim to infer latent residue–atom correspondences [27–29], yet, whether these attention patterns reflect true physical interactions has never been systematically evaluated. A fundamental question therefore remains unanswered: do learned attention representations capture the non-covalent forces that actually govern binding, or merely correlate with binding likelihood?

To address this challenge, here we present **ExplainBind**, a protein–ligand binding framework that jointly predicts binding likelihood, localizes binding sites at the residue level, and decodes the underlying non-covalent interaction patterns from sequence alone, without requiring three-dimensional structural inputs. Binding can be represented at multiple scales: from the full binding complex and pocket to a residue-level interaction profile and finally to interaction-type–specific binding sites (Fig. 1a). To enable training and evaluation at the residue and interaction-level, we construct **InteractBind**, a curated protein–ligand benchmark derived from experimentally resolved complexes and annotated with residue–atom interaction maps. By explicitly supervising its cross-attention module with physically grounded interaction maps, ExplainBind bridges predictive accuracy and mechanistic interpretability, moving beyond the binary binding classifiers that currently dominate the field (Fig. 1b,c). Evaluated across a controlled similarity spectrum from closely related to deliberately dissimilar protein and ligand sequences, ExplainBind achieves state-of-the-art performance while its learned attention maps reliably predict binding-site localization. Applied retrospectively to angiotensin-converting enzyme (ACE) to enrich potent inhibitors, and prospectively to L-2-hydroxyglutarate dehydrogenase (L2HGDH) to distinguish inhibitors from activators, the framework demonstrates broad utility for mechanistically informed drug discovery and enzyme engineering.

**Figure 1:**
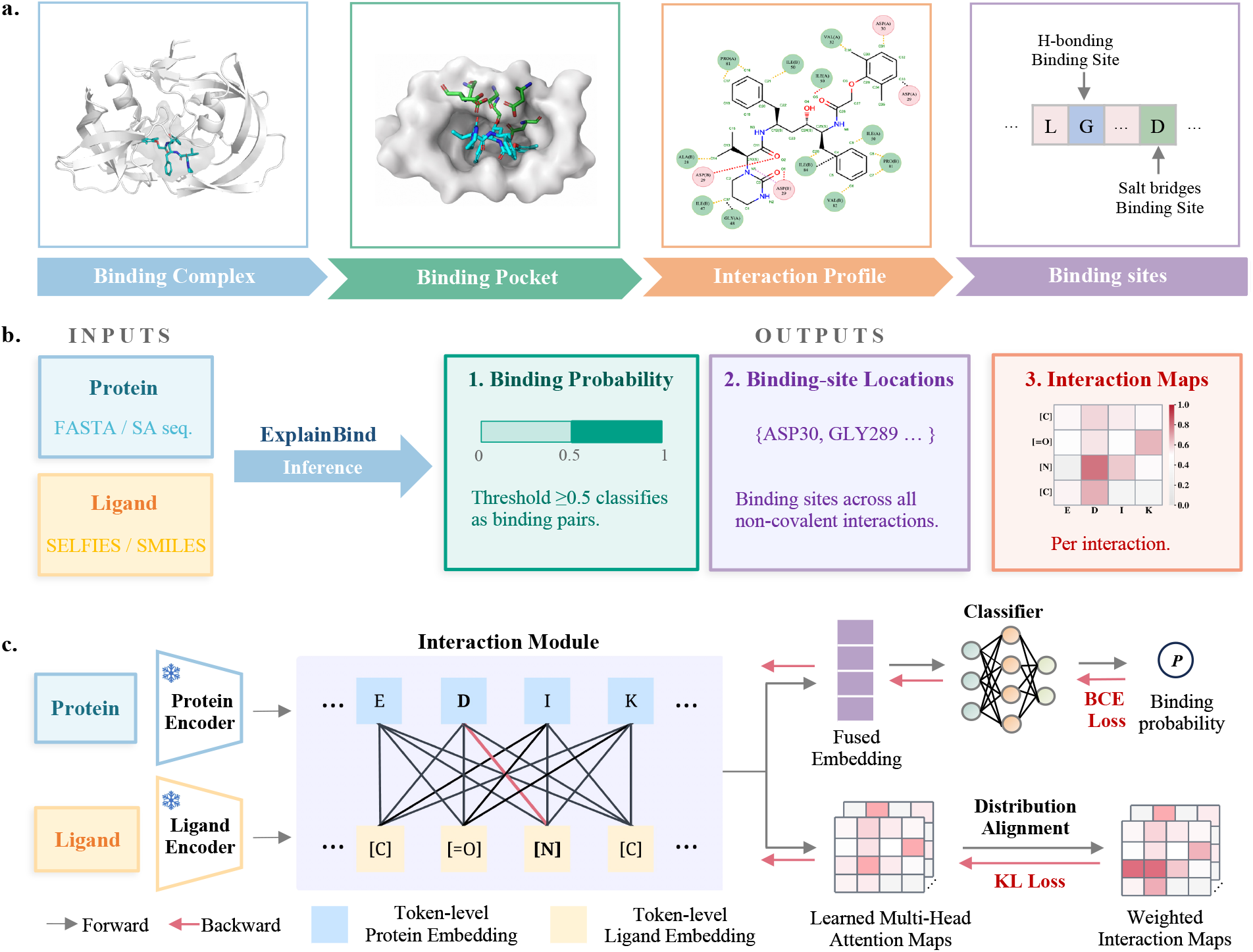
Multi-scale protein–ligand binding representation and ExplainBind framework. **a**. Multi-scale representation of protein–ligand binding: binding complex, binding pocket, interaction profile, and interaction-type–specific binding sites. **b**. During inference, ExplainBind takes a protein sequence and ligand representation as input and predicts binding probability, localizes binding-site residues, and generates interaction maps across six non-covalent interaction types. **c**. During training, frozen protein and ligand encoders produce token-level embeddings fused by an interaction module, with a BCE loss supervising binding prediction and a weighted KL divergence loss grounding multi-head attention maps in ground-truth interaction maps from InteractBind.

## 2 Results

### 2.1 Overview of ExplainBind

**ExplainBind** is an end-to-end framework that connects protein–ligand binding prediction with binding-site localization and mechanistic modeling of the physicochemical interactions that mediate molecular recognition. As shown in Fig. 1c, the framework uses frozen foundation encoders with an interaction module to capture token-level interactions between amino acid residues and ligand atoms, supervised using interaction-type–specific maps derived from curated protein–ligand complexes across six non-covalent interactions. Our proposed framework is jointly optimized with two complementary objectives: a Binary Cross-Entropy (BCE) loss [30] on the predicted binding probability and a Kullback–Leibler divergence (KL) loss [31] aligning the learned attention maps with ground-truth non-covalent interaction maps, grounding predictions in biologically meaningful residue–atom relationships.

Underpinning this framework is **InteractBind**, a novel large-scale database that we constructed by systematically mining and annotating protein–ligand complexes from the Protein Data Bank (PDB) [32]. We localize the binding pocket from each complex and then extract non-covalent interaction annotations. Non-covalent interactions are identified using rule-based criteria grounded in geometric and chemical constraints, with interaction-specific distance cutoffs and optional angular filters applied to ensure validity (Supplementary Table S1). Interaction strength is modelled by a distance-dependent piecewise linear decay function that maps interatomic or geometric distances within the cutoff window to continuous values in [10^*−*6^, 1], yielding an *m × n* interaction map per interaction type, where *m* and *n* denote the numbers of ligand units and protein residues, respectively. In addition, The six non-covalent interaction maps are aggregated into an overall interaction map. Finally, residue-level binding-site labels are obtained by projecting the overall interaction map onto the residue dimension.

We evaluate ExplainBind and seven representative baselines (Supplementary S.2.4) under both in-distribution (**ID**) and out-of-distribution (**OOD**) settings. For ID, we use the affinity-aware subset from InteractBind, stratifying pairs according to binding affinity (kcal/mol) and retaining only high-confidence negative samples (*≥ −*5 kcal/mol) and positive samples (*≤ −*7 kcal/mol) for supervision, whilst excluding cases of moderate affinity to reduce label ambiguity. For the OOD setting, we construct similarity-controlled evaluation scenarios along two complementary axes. Ligand chemical similarity is quantified using Tanimoto similarity over extended connectivity fingerprints (ECFP) [33], yielding mean train–test similarities of 8%, 35%, 40%, and 59%, with protein similarity fixed at 33% across all splits. Protein similarity is quantified using global sequence alignment based on the Needleman–Wunsch algorithm [34], yielding peak train–test similarities of 25%, 28%, 31%, and 33%. Combining these regimes results in eight OOD datasets spanning progressively increasing degrees of chemical and sequence novelty. For binary binding prediction, model performance is assessed using accuracy (Acc), area under the receiver operating characteristic curve (AUROC), area under the precision–recall curve (AUPRC), and F1 score. For binding-site localization, we report Binding Residue Hit Rate (BRHR) and Recall, whereas for non-covalent interaction prediction, we report Interaction Hit Rate (IHR) and Recall, following the evaluation protocol defined in Supplementary S.2.3.

### 2.2 From binary binding prediction to binding-site localization

We evaluated ExplainBind and seven representative baselines on both binary binding prediction and binding-site localization, asking whether the strong classification performance characteristic of current sequence-based methods translates into accurate identification of the non-covalent contacts that mediate protein–ligand recognition. The answer is an unambiguous no. Across seven representative baselines spanning from cross-attention architectures (MolTrans [35], TransformerCPI [20], CAT-DTI [27], HyperAttentionDTI [21], PerceiverCPI [36]) to bilinear attention mechanisms (DrugBAN [28], GraphBAN [37]), binary binding prediction approaches saturation under in-distribution conditions, with AUROC ranging from 95.1% to 97.8% and accuracy from 84.1% to 89.3% (Table 1). Yet, binding-site localization remains uniformly and severely limited: BRHR@1 ranges from only 10.6% to 16.1% across all baselines, meaning that even the strongest model fails to recover binding residues in its top prediction for more than 83% of protein–ligand pairs.

**Table 1:** Performance (%) comparison with baselines under the in-distribution setting. Results are mean*±* standard deviation over 5-fold cross-validation. BRHR@*K* measures whether at least one of the Top-*K* highlighted protein residues matches a ground-truth binding residue. The best results are shown in **bold**, and the second-best results are underlined.

| Model | Binary Binding Prediction |  |  |  | Binding-site Localization |  |  |  |  |
| --- | --- | --- | --- | --- | --- | --- | --- | --- | --- |
|  | ACC | AUPRC | AUROC | F1 | BRHR@1 | BRHR@3 | BRHR@5 | Recall@5 | Recall@10 |
| MolTrans | 88.6 ± 0.7 | 93.7 ± 1.2 | 95.1 ± 2.2 | 88.1 ± 0.5 | 10.9 ± 0.5 | 17.7 ± 0.2 | 22.4 ± 0.8 | 1.7 ± 0.4 | 3.5 ± 0.3 |
| TransformerCPI | 89.1 ± 0.6 | 90.8 ± 0.8 | 95.4 ± 0.3 | 88.5 ± 0.6 | 10.9 ± 0.2 | 16.7 ± 0.7 | 21.7 ± 0.4 | 1.7 ± 0.3 | 3.4 ± 0.4 |
| HyperAttentionDTI | 84.1 ± 0.9 | 92.8 ± 0.3 | 95.5 ± 0.6 | 88.5 ± 0.5 | 12.3 ± 0.4 | 17.7 ± 0.6 | 20.8 ± 0.3 | 1.8 ± 0.3 | 3.9 ± 0.2 |
| PerceiverCPI | 86.6 ± 0.8 | 92.6 ± 0.5 | 96.9 ± 0.2 | 82.2 ± 0.7 | 10.6 ± 0.1 | 13.9 ± 0.5 | 19.9 ± 0.9 | 1.6 ± 0.2 | 3.1 ± 0.2 |
| CAT-DTI | 88.1 ± 0.5 | 92.5 ± 0.6 | 96.8 ± 0.4 | 87.0 ± 0.3 | 12.2 ± 0.7 | 14.8 ± 0.2 | 22.1 ± 0.6 | 1.2 ± 0.2 | 2.8 ± 0.3 |
| DrugBAN | 89.3 ± 0.4 | 94.3 ± 0.4 | 97.6 ± 1.2 | 87.5 ± 1.4 | 15.7 ± 0.6 | <u>23.9 ± 0.1</u> | <u>31.4 ± 0.4</u> | <u>2.1 ± 0.2</u> | 3.7 ± 0.1 |
| GraphBAN | 88.9 ± 0.3 | <u>94.4 ± 0.3</u> | <u>97.8 ± 0.2</u> | <u>88.5 ± 0.4</u> | <u>16.1 ± 0.5</u> | 22.9 ± 0.2 | 30.4 ± 0.6 | 1.9 ± 0.2 | <u>4.1 ± 0.3</u> |
| **ExplainBind** | **91.2 ± 0.4** | **95.4 ± 0.2** | **99.3 ± 0.3** | **90.5 ± 0.6** | **55.6 ± 0.7** | **69.5 ± 0.2** | **74.6 ± 0.6** | **7.3 ± 0.3** | **8.7 ± 0.2** |

Critically, high binary classification performance provides no reliable indication of localization ability. PerceiverCPI achieves a strong AUROC of 96.9% but the lowest BRHR@1 of 10.6%, while GraphBAN obtains the strongest baseline AUROC and F1 yet is surpassed by DrugBAN on BRHR@3, BRHR@5, and Recall@5. These crossovers suggest that binding prediction and binding-site localization remain decoupled under binding label-only supervision. By explicitly supervising cross-attention with physically grounded, interaction-type-specific maps, ExplainBind achieves 55.6% BRHR@1, 69.5% BRHR@3, and 74.6% BRHR@5, improving over the strongest baselines by 39.5, 45.6, and 43.2 percentage points, respectively. It also improves Recall@5 and Recall@10 to 7.3% and 8.7%, while achieving the best binary prediction performance across all metrics: 91.2% accuracy, 95.4% AUPRC, 99.3% AUROC, and 90.5% F1. Additional ablation studies analysing the contribution of each component are provided in Supplementary S.3.8.

### 2.3 Generalization under similarity-controlled evaluation

Generalization of sequence-based binding models is further evaluated under carefully controlled out-of-distribution settings based on protein and ligand similarity. In the protein-controlled setting, test proteins are constrained to have limited sequence similarity to training proteins, modeling generalization to unseen target families (Fig. 2a). Across models, performance decreased as the train–test sequence-similarity threshold became more stringent, indicating that protein sequence divergence is a major source of distribution shift. This trend suggests that conventional in-distribution evaluation can overestimate performance when sequence-based binding models are applied to novel protein targets. Despite this challenging setting, ExplainBind achieves the strongest performance, with an AUROC of 85 at the most stringent 25% threshold and 94 at 33%, showing greater robustness to target-side novelty than existing baselines.

**Figure 2:**
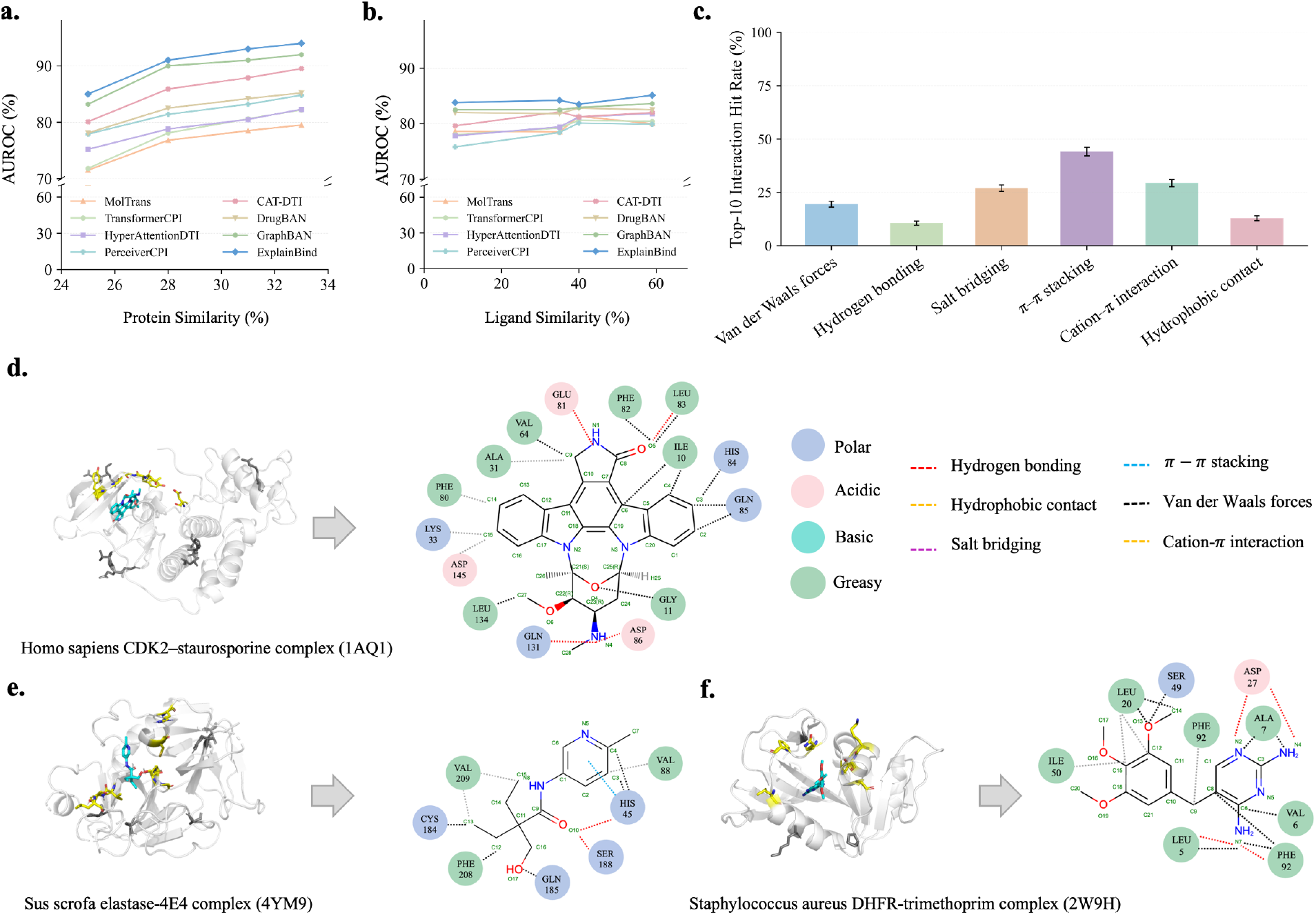
Out-of-distribution binding prediction and non-covalent interaction prediction via ExplainBind. **a**. OOD performance under protein sequence similarity–controlled splits, with AUROC increasing consistently as sequence similarity increases. **b**. OOD performance under ligand similarity–controlled splits, showing stable AUROC across ligand-similarity thresholds. **c**. Interaction-specific non-covalent interaction prediction measured by Top-10 Interaction Hit Rate (IHR@10) across non-covalent interaction types. **d-f**. Case studies of representative protein–ligand complexes (PDB: 1AQ1, 4YM9 and 2W9H). Interaction maps derived from crystal structures are compared with model predictions. The top 20 predicted residues recover all experimentally observed contacts. Ligands are shown in cyan, the top 1 predicted residue is highlighted in yellow and lies within the experimental binding site, whereas lower-ranked residues among the top 20 that fall outside the annotated site are shown in black.

For ligand-controlled splits, test ligands are constrained to be chemically dissimilar from the training ligands (Fig. 2b). In contrast to the protein-controlled setting, all models show a narrower performance range across ligand similarity thresholds, suggesting that chemical diversity among ligands produces a comparatively modest distribution shift when protein targets are sufficiently represented. ExplainBind maintains stable performance, with AUROC values ranging from 83 to 85 across ligand similarity thresholds from 8% to 59%, and remains consistently competitive with the baselines. Together, these results show that target-side novelty is the dominant challenge for sequence-based binding prediction, while ExplainBind provides stronger OOD generalization across protein- and ligand-controlled screening scenarios.

### 2.4 Prediction of non-covalent interactions

Beyond binding-site localization, ExplainBind resolves the specific non-covalent forces mediating each residue–atom contact, producing six interaction-type-specific maps for each protein–ligand pair (Fig. 2c). To evaluate its performance, we apply a strict exact-match criterion using the Top-10 Interaction Hit Rate (IHR@10), where at least one of the ten highest-scoring residue–atom pairs must match a true contact among approximately 12,880 candidates (368 protein residues *×* 35 ligand atoms), of which fewer than three are annotated contacts on average, giving a random success rate of approximately 0.23%. Despite this stringency, ExplainBind recovers ground-truth contacts across all six interaction types. *π*–*π* stacking achieves the highest IHR@10 (44.2%), consistent with the distinctive aromatic residue signatures that mediate this interaction. Cation–*π* interactions and salt bridges also show strong performance, reaching 29.4% and 27.0%, respectively, reflecting the chemically constrained residue preferences associated with charged and aromatic contacts. Van der Waals forces achieve 19.5%, whereas hydrophobic contacts (12.9%) and hydrogen bonding (10.6%) are more challenging, likely because they involve broader residue and atom types and are less defined by simple sequence-level chemical signatures. Detailed interaction-specific IHR@*K* and Recall@*K* results, together with their analysis, are provided in Supplementary S.3.1.

### 2.5 Case studies of non-covalent interaction recognition

We examined three co-crystallized protein–ligand complexes from distinct biological contexts to assess whether ExplainBind’s predicted interaction maps agree with crystallographically resolved contacts: *Homo sapiens* CDK2–staurosporine, *Sus scrofa* elastase–4E4, and *Staphylococcus aureus* DHFR–trimethoprim (Fig. 2d-f). For CDK2–staurosporine, ExplainBind recovered hydrogen-bonding residues including GLU81, LEU83, ASP86, and GLN131, alongside hydrophobic and van der Waals contacts that stabilize the ligand within the kinase binding cleft. For elastase–4E4, the model identified hydrogen bonds near the catalytic triad, hydrophobic contacts around VAL88, VAL209, and PHE208, and a *π*–*π* stacking interaction involving HIS45. For DHFR– trimethoprim, ExplainBind captured the mixed polar–apolar active-site environment, including hydrogen bonds at LEU5, ASP27, and PHE92, and hydrophobic contacts at ILE50, LEU20, and PHE92. Across all three complexes, the predicted interaction patterns agreed with crystallographically observed contacts, demonstrating that ExplainBind recovers chemically interpretable recognition features without structural input. Such outputs provide experimentally actionable hypotheses for identifying ligand-recognition residues, prioritizing mutagenesis, and guiding mechanism-aware ligand optimization (Supplementary S.3.2).

### 2.6 Prioritization of high-activity ligands

To evaluate ExplainBind under a ligand prioritization setting, we performed retrospective enrichment analysis on angiotensin-converting enzyme (ACE), a key regulator of blood pressure and cardiovascular homeostasis [38], and wet-lab validation on L-2-hydroxyglutarate dehydrogenase (L2HGDH), a mitochondrial metabolic enzyme for which both activation and inhibition are of potential therapeutic interest [39]. Neither target appears in the InteractBind training set or has a closely related homolog in the training data, providing a stringent evaluation on genuinely unseen proteins.

#### Retrospective prioritization on ACE

As shown in Fig. 3a, ACE ligands span a broad range of IC_50_ affinity strata. When ranked by predicted binding probability, ultra-potent ligands (IC_50_ *≤* 1 nM) are preferentially enriched among the top-ranked candidates, with enrichment decreasing smoothly as the ranking cutoff Top-*K* increases from 25 to 200 (Fig. 3b). The Top-25 ranked compounds achieve a hit rate of 0.16, demonstrating that highly potent ligands are preferentially concentrated among top-ranked candidates. Consistent with this trend, mean predicted binding probability decreases with increasing Top-*K* when compounds are ordered by experimental activity, decreasing from 0.486 at Top-20 to approximately 0.293 at Top-200 (Fig. 3c).

**Figure 3:**
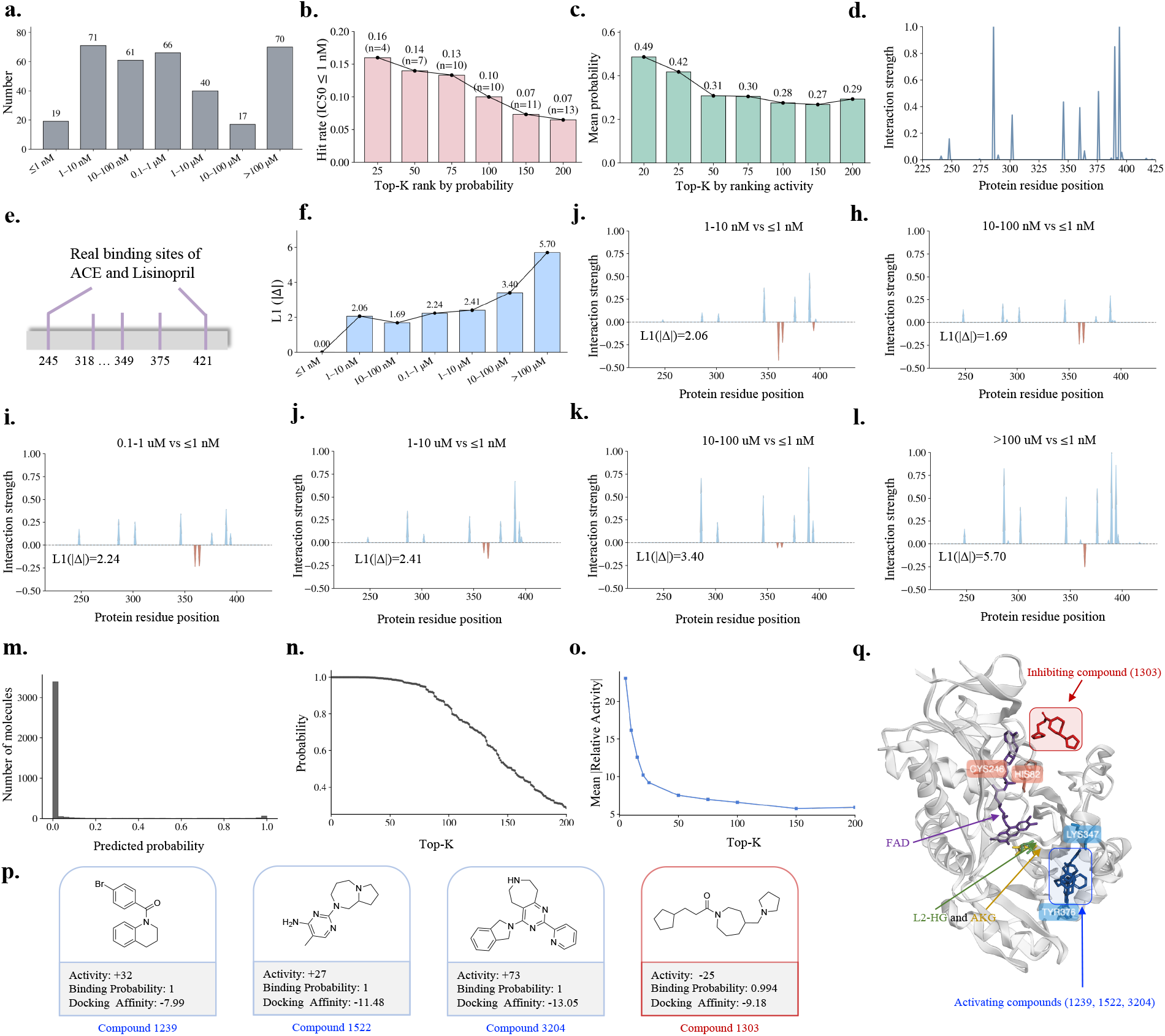
Ligand prioritization for ACE and L2HGDH. **a**. Distribution of ACE ligands across IC_50_ affinity strata. **b**. Enrichment of highly potent compounds (IC_50_ ≤1 nM) among the Top-*K* candidates ranked by predicted binding probability. **c**. Mean predicted binding probability across Top-*K* candidates ranked by experimental activity. **d**. Predicted ACE binding-site interaction landscape. **e**. Experimentally resolved ACE–lisinopril binding sites. **f**. Overall shift in predicted interaction patterns, measured by *𝓁*_1_(| Δ|), between each lower-affinity stratum and the IC_50_ ≤ 1 nM reference group. **g–l**. Position-wise interaction changes (Δ) across affinity strata relative to the IC_50_ ≤1 nM group, with positive and negative values denoting enhanced and reduced predicted interactions, respectively. **m**. Distribution of predicted binding probabilities across the internal compound pool (3,750 compounds) with L2HGDH, broadly consistent with the experimentally observed activity spectrum. **n**. The binding probability ranking exhibits a rapid decay from 1 to 0.3 for the top 200 compounds with L2HGDH. **o**. The mean |Relative Activity| within the top 200 compounds indicates a significant enrichment of active compounds among the highly ranked candidates. **p**. The highly active ligands within the Top-50 compounds, illustrating scaffold diversity among both experimentally validated inhibitors and activators. **q**. Mechanistic basis of L2HGDH modulation by representative ligands. Docked poses of activators (compounds 1239, 1522, and 3204; blue) and inhibitor (compound 1303; red) superimposed on the L2HGDH structure (PDB: 8W78, chain A; semi-transparent surface), as determined by global docking. The natural substrate L-2-hydroxyglutarate (L2-HG; green), product *α*-ketoglutarate (AKG; yellow), and FAD cofactor (purple) are shown for reference. ExplainBind-predicted hydrogen-bonding interactions reveal mechanistically distinct binding modes: inhibitor engagement is mediated by Cys246 and His82, residues situated in the vicinity of the FAD-binding region, suggesting that inhibitor occupancy at this site may sterically compress the FAD cofactor deeper into the catalytic pocket, occluding substrate entry and thereby abolishing catalytic activity. In contrast, activator binding is coordinated through Lys347 and Tyr376, which are spatially distinct from the FAD pocket.

**Figure 4:**
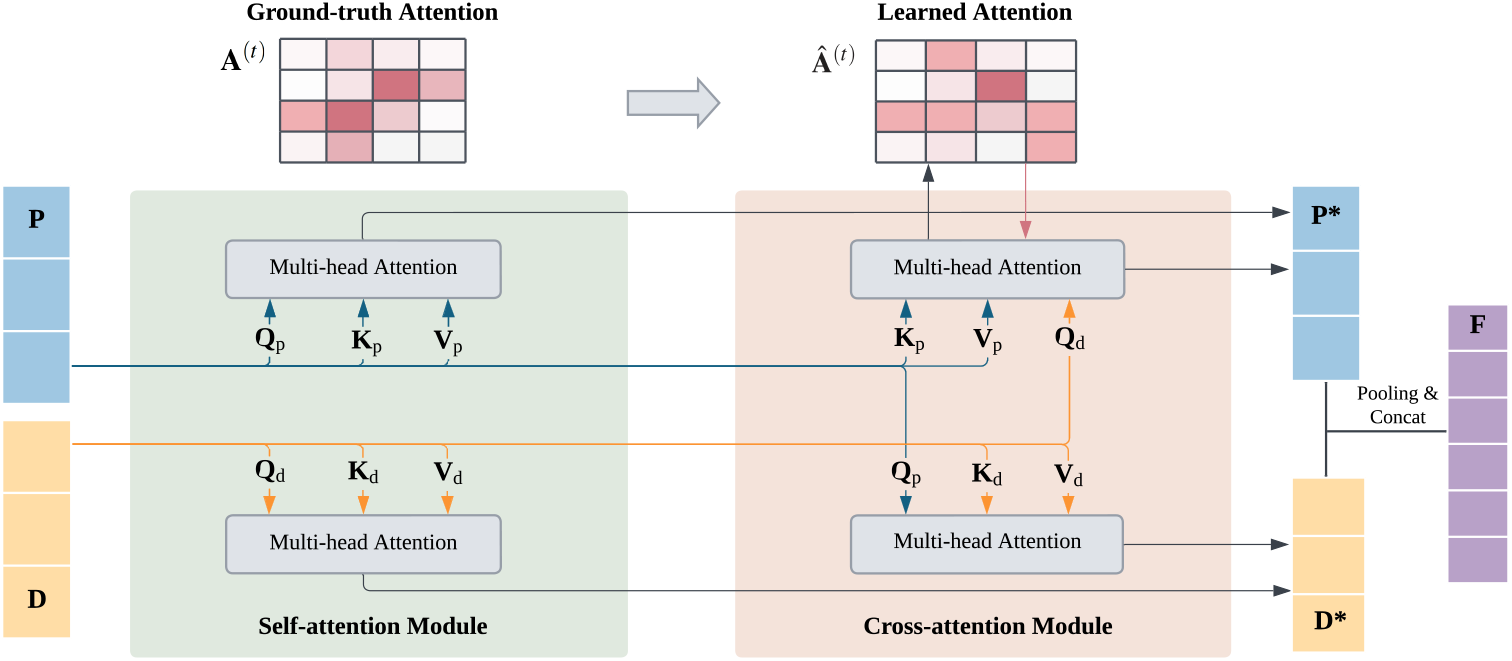
Interaction Module: Token-level embeddings of proteins (**P**) and ligands (**D**) are refined by self-attention and aligned by cross-attention. Multi-head attention yields per-head interaction maps {**Â** ^(*t*)^}, which can be supervised against ground-truth maps {**A**^(*t*)^} when available. Fused embeddings **P**^*\**^ and **D**^*\**^ are mean-pooled and concatenated into **F**.

Beyond compound-level ranking, ExplainBind provides interaction profiles that reveal affinity-dependent patterns of molecular recognition. The predicted ACE interaction landscape is broadly consistent with experimentally resolved ACE–lisinopril binding sites [40] (Fig. 3d,e). Comparing interaction patterns across progressively weaker affinity strata relative to the IC_50_ *≤* 1 nM reference group (Fig. 3f–l), interaction differences remain localised and modest at nanomolar potency (*𝓁*_1_ norms: 1.687–2.236), but become increasingly pronounced as affinity decreases into the micromolar regime (*𝓁*_1_ norm: 3.400 for 10–100 *µ*M; 5.703 for IC_50_ *>* 100 *µ*M), indicating systematic erosion of key interaction patterns with reduced binding potency.

#### Wet-lab validation on L2HGDH

ExplainBind is applied to an internal pool of 3,750 compounds to predict binding probability and guide experimental selection for this mitochondrial enzyme active in central carbon metabolism [41]. The majority of compounds are assigned low predicted probabilities below 0.2 (Fig. 3m), with the Top-200 spanning from 1.0 to approximately 0.3 (Fig. 3n), motivating their selection for experimental testing, where relative activity was measured as the percent change in enzyme activity relative to vehicle control (Supplementary S.2.7). Among experimentally tested compounds, the Top-5 ranked achieve a mean |Relative Activity| of 22.40, which decreases rapidly to 9.65 at Top-20 and reaches a shallow tail of approximately 5–7 by Top-200 (Fig. 3o), consistent with progressive inclusion of weaker modulators and a coherent transition from high-confidence, high-activity candidates to lower-confidence predictions.

Experimental measurements confirm strong enrichment of functional modulators among top-ranked compounds. Among the Top-50, four compounds exhibit high experimental activity (|Relative Activity| *≥* 25), including three activators and one inhibitor (Fig. 3p), all assigned high predicted binding probabilities. Global docking of these four compounds against L2HGDH (PDB: 8W78, chain A) is performed to characterize their binding poses, and as these ligands are potent hydrogen-bond acceptors, ExplainBind-predicted hydrogen-bonding interactions are further interrogated to elucidate their mechanistically distinct binding modes: inhibitor engagement (compound 1303) is mediated by Cys246 and His82, residues situated in the vicinity of the FAD-binding region, suggesting that inhibitor occupancy at this site sterically compresses the FAD cofactor deeper into the catalytic pocket, occluding substrate entry and thereby abolishing catalytic activity; whereas activator binding (compounds 1239, 1522, and 3204) is coordinated through Lys347 and Tyr376, residues spatially distinct from the FAD pocket, potentially contributing to an allosteric mechanism of activation (Fig. 3q).

## 3 Discussion

Mechanistic insight into molecular recognition has historically been considered inseparable from three-dimensional structural data—an assumption that has long restricted sequence-based approaches to binary binding predictions of whether binding occurs, leaving the spatial (“where”) and mechanistic (“how”) dimensions of interaction entirely unresolved. ExplainBind fundamentally challenges this structural dogmatism by demonstrating that the essential physical determinants of molecular recognition are recoverable from sequence, provided that the underlying deep-learning architecture is conditioned not merely on binding labels but also on the explicit non-covalent interaction topologies that physically mediate the binding event. The indispensable catalyst for this predictive capacity is not explicit atomic geometry, but rigorous biophysical grounding.

Consequently, our most important finding is not simply that ExplainBind predicts binding more accurately, but that it resolves where and how binding occurs with high resolution—a distinction that yields immediate translational utililty. Delineating the exact residues that anchor a ligand and through which non-covalent forces establishes a useful framework rational mutagenesis, enables precise selectivity engineering, and constrains the chemical space of productive modifications in ways that a monolithic binding probability alone cannot. For ACE, this translates into affinity-stratified interaction landscapes that systematically identify which contacts are dispensable as binding potency decays. This provides directly actionable structural insights lead optimization in the complete absence of empirical or modeled 3D structural data. For L2HGDH, a target entirely absent from training, ExplainBind successfully differentiated inhibitors from activators through differential interaction profiles anchored to mechanistically distinct residues: inhibitor engagement was localized in close proximity to the FAD-binding region, strongly implying a mechanism of competitive steric occlusion that blocks substrate entry. Conversely, activator coordination was mapped to spatially segregated residues, pointing toward a positive allosteric modulation mechanism—a resolution fundamentally inaccessible to conventional models trained on binary binding labels alone. That this precise discrimination was achieved prospectively, on a genuinely unseen target, underscores that ExplainBind’s interaction profiles carry functional predictive power that extends far beyond simple binding likelihoods, offering profound implications for context-dependent therapeutic targeting of metabolic enzymes.

Sequence-based protein–ligand models have increasingly adopted cross-attention mechanisms with the implicit expectation that meaningful residue–atom correspondences would spontaneously from binding label supervision alone. Whether the resulting attention patterns reflect the actual non-covalent forces that govern molecular recognition, however, has never been systematically tested. ExplainBind addresses this challenge directly, showing that attention maps grounded in experimentally derived interaction profiles successfully recover contact patterns that mirror crystallographically resolved binding interfaces across chemically and phylogenetically diverse complexes. This finding sharply distinguishes the ExplainBind from end-to-end structure-based co-folding models such as AF3, RoseTTAFold All-Atom, Chai-1 and Boltz-1. While these deep-learning co-folding models generate plausible binding geometries, they frequently yield local coordinates that violate fundamental steric and electrostatic constraints, resulting in atomic clashes or unphysical ligand distortions [19]. This divergence suggests that geometric fidelity and chemical realism are dissociable, and that capturing the latter mandates explicit physicochemical conditioning and supervision. More broadly, any model trained solely on binary binding outcomes will inevitably converge on statistical correlates of binding rather than its causal physical drivers. This critical distinction becomes the limiting bottleneck precisely when translating a model’s utility from retrospective classification tasks to prospective, mechanistically sound hypothesis generation.

The convergence of scalable, sequence-based inference with mechanistic transparency positions ExplainBind as a versatile, broadly applicable tool for contexts where structural data are absent, low-resolution, or entirely unresolvable. These applications range from high-throughput ligand prioritization across understudied protein families to the rational engineering of industrial or therapeutic enzymes with reprogrammed substrate specificities. Looking forward, extending interaction-grounded supervision to multi-target selectivity profiling and integration with generative molecular design frameworks, where interaction-type constraints could guide scaffold optimization toward desired binding modes, represents a natural and immediate direction. As experimentally annotated interaction data continue to expand through structural genomics and screening efforts, frameworks that ground and anchor learned representations in foundational physical interaction principles will become increasingly central to predictive molecular biology.

## 4 Methods

### 4.1 Problem Formulation

We formulate protein–ligand binding (PLB) modeling as a joint learning problem over three complementary tasks: (1) binary binding prediction, (2) binding-site localization, and (3) non-covalent interaction prediction. These tasks share the same protein–ligand input but differ in their prediction objectives.

#### Binary binding prediction

Given a protein sequence *𝒫* = (*p*_1_, …, *p*_*n*_) composed of residue-level tokens and a ligand sequence *𝒟* = (*d*_1_, …, *d*_*m*_) representing atom-level structure, the objective of binary binding prediction is to determine whether the protein–ligand pair forms a binding interaction. Formally, this task can be written as

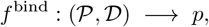

where *p* ∈ [0, 1] is a scalar probability indicating the likelihood that the protein–ligand pair interacts.

#### Binding-site prediction

In addition to binary binding prediction, PLB modeling requires localization of the protein residues and ligand atoms involved in binding. Given the same protein sequence *𝒫* and ligand sequence *𝒟*, the objective is to identify the set of protein residues and ligand atoms that participate in the binding mechanism. Formally, this task is defined as

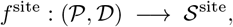

where *𝒮*^site^ = *{*(*d*_*i*_, *p*_*j*_)*}* denotes the predicted list of ligand-atom and protein-residue pairs involved in binding. Each pair (*d*_*i*_, *p*_*j*_) ∈ *𝒮*^site^ indicates that ligand atom *d*_*i*_ and protein residue *p*_*j*_ are predicted to participate in the binding.

#### Non-covalent interaction prediction

Beyond identifying binding-site locations, PLB modeling further requires distinguishing the physicochemical interaction types that mediate protein–ligand recognition. Given the same protein sequence *𝒫* and ligand sequence *𝒟*, the objective is to predict a set of interaction-specific residue–atom maps, each corresponding to a particular class of non-covalent interaction. Formally, this task is defined as

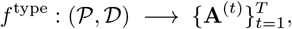

where *T* denotes the number of non-covalent interaction types. Each matrix **A**^(*t*)^ *∈* ℝ^*m×n*^ represents the residue–atom interaction pattern associated with non-covalent interaction type *t*, and each entry 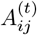indicates the predicted likelihood that ligand atom *d*_*i*_ and protein residue *p*_*j*_ form interaction type *t*.

Together, these tasks characterize protein–ligand binding at increasing levels of resolution: whether binding occurs, where the binding occurs, and which physicochemical interactions mediate recognition.

### 4.2 Model Architecture

#### 4.2.1 Encoders

Capturing rich and biologically grounded representations is essential for exploring fine-grained and explainable protein–ligand interactions. For ligands, SMILES strings are widely used but may contain invalid segments and lose critical structural cues [42]. To overcome these limitations, we adopt SELFIES [42], a robust molecular grammar that guarantees valid molecular graphs and preserves atom-level information [42]. For proteins, we primarily use FASTA sequences, in which each token corresponds to one of the 23 standard amino acid symbols. When reliable structural information is available, ExplainBind can optionally incorporate structure-aware (SA) protein representations derived via *Foldseek* [43], providing explicit local structural context [44].

Foundation models have shown strong capability in encoding contextual and structural information through large-scale pre-training on biological corpora. The framework combines frozen foundation encoders for proteins and ligands, enabling ExplainBind to leverage powerful pretrained representations without updating the backbone models. In our default configuration, we use ESM-2 [45] as the frozen protein encoder and SELFormer [46] as the frozen ligand encoder for SELFIES representations. These encoders produce token-level protein embeddings **P** and ligand embeddings **D** that serve as inputs to the interaction module, forming the basis for accurate binding prediction and explainable interaction modeling, including SaProt [44] when SA representations are used, as well as ProteinBERT [47], MoLFormer [48], and ChemBERTa-2 [49], with results summarized in Supplementary S9.

#### 4.2.2 Interaction module

Unlike prior unsupervised models for PLB prediction, ExplainBind explicitly supervises attention with ground-truth maps derived from experimentally resolved protein–ligand complexes. Concretely, we convert residue–level contacts in each complex into per-type interaction maps and use them to directly optimize the attention computed from protein and ligand embeddings. To our knowledge, our proposed framework is the first to use complex-level attention maps to guide token-level fusion of proteins and ligands. Unsupervised attention conflates heterogeneous physical forces and tends to produce diffuse, non-identifiable alignments. By supervising attention with per-type contact maps, we (a) disambiguate interaction mechanisms (e.g., hydrogen bond vs. hydrophobic contact), (b) improve the faithfulness of explanations, and (c) use the same supervision signal to shape the fused token embeddings towards binding-relevant substructures.

Let **D** ∈ ℝ^*m×h*^ and **P** ∈ ℝ^*n×h*^ denote the token-level embeddings of the ligand and protein, respectively. Linear projections generate query (**Q**), key (**K**), and value (**V**) matrices for each modality:

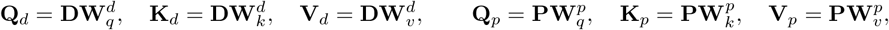

where 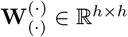are learnable projections.

The interaction module uses multi-head attention with *H* = 8 heads: six heads correspond to non-covalent interactions, and two heads capture overall binding patterns. For each head *t ∈ 𝒯*, bidirectional cross-attention computes the residue–level interaction intensity (where *h* is the hidden dimension size):

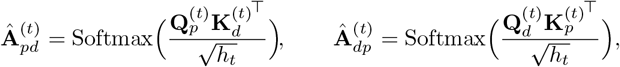

where 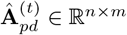and 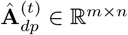. Self-attention is simultaneously applied to refine intra-sequence contexts:

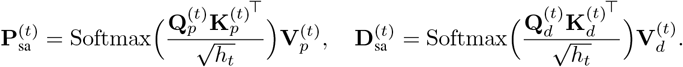

Type-specific fused representations are obtained by combining self- and cross-attention outputs:

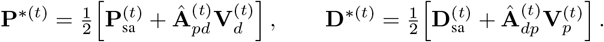

Finally, outputs from all heads are concatenated and projected to yield the fused token sequences:

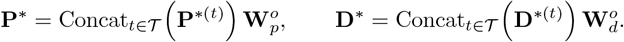

#### 4.2.3 Classifier

Given the fused token embeddings **P**^*\**^ ∈ ℝ^*n×d*^ and **D**^*\**^ ∈ ℝ^*m×d*^, we form a pair-level representation by mean pooling over tokens and concatenating the pooled vectors:

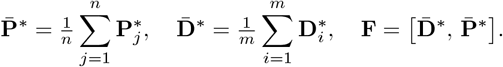

The binding probability is predicted by a lightweight MLP head, *p* = MLP(**F**).

#### 4.2.4 Training objective

Let *y ∈* {0, 1} denote the ground-truth binding label and *p* the predicted binding probability. The model is optimized using a binary cross-entropy classification loss,

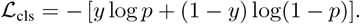

When interaction supervision is available, we further align the predicted cross-attention maps with ground-truth interaction annotations. For each supervised head *t* ∈ *𝒯*_sup_, the predicted attention **Â** ^(*t*)^ and the ground-truth interaction map **A**^(*t*)^ are normalised over residue–atom pairs to form discrete distributions, 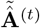and **Ã** ^(*t*)^, respectively. The attention alignment loss is defined as a weighted KL divergence,

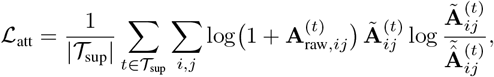

where 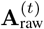denotes the unnormalised interaction intensity used to construct **A**^(*t*)^. For training pairs without interaction annotations, only *ℒ*_cls_ is applied.

The final objective is

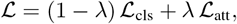

where *λ ∈* [0, 1] controls the contribution of attention supervision. In practice, *λ* is selected based on validation performance via an ablation study, and we set *λ* = 0.3 for all experiments, which achieves the best overall trade-off between predictive accuracy and attention alignment (see Supplementary S.3.8).

## 5 Data and code availability

The InteractBind database is derived from publicly available protein–ligand complex structures deposited in the Protein Data Bank (PDB). **InteractBind has been uploaded to the manuscript submission system for peer review**. The source code required to reproduce the results of this study is publicly available at **GitHub**: https://github.com/ZhaohanM/ExplainBind. A user-friendly web interface is accessible via the **Demo UI**: https://huggingface.co/spaces/Zhaohan-Meng/ExplainBind.

## 6 Acknowledgments

This work is supported in part by National Institutes of Health grants HL155107 and HL166137, and by American Heart Association MERIT award AHA1185447 to J.L. This work is also supported in part by an Ignite Grant from Brigham and Women’s Hospital to W.O. and J.L. H.X. acknowledges the U.S. National Science Foundation’s national research cyberinfrastructure ecosystem through ACCESS under award number CIS250621. K.Y. acknowledges support from Cancer Research UK (EDDPGM-Nov21/100001, DRCMDP-Nov23/100010 and core funding to the CRUK Scotland Institute (A31287)), BBSRC BB/V016067/1, Prostate Cancer UK MA-TIA22-001 and EU Horizon 2020 grant ID: 101016851. We thank Christy Biji, Christian K. Soule, Brittany A. Marion, and Jane K. McIninch (Center for the Development of Therapeutics, The Broad Institute of MIT and Harvard) for their support in the wet-lab screening experiments.

## 7 Author contributions

Z.M. (Zaiqiao Meng), H.X. and J.L. conceived the study and provided overall supervision. Z.M. (Zhaohan Meng) designed the study, developed the model, performed the experiments, analyzed and interpreted the results, and prepared the manuscript. Z.B. constructed the dataset, performed data analysis, and prepared the manuscript. I.O., K.Y., J.S. and Y.Z. provided additional supervision and domain expertise. W.O., J.C., W.J., A.S., and K.L. led the wet-lab validation. All authors contributed to the writing, reviewed the manuscript, and approved the final version.

## Supplementary Material

### S.1 Preliminary

#### S.1.1 Covalent bonds and non-covalent interactions

Molecular systems involve two fundamental types of interactions: <u>covalent bonds</u> and <u>non-covalent interactions</u> [4]. Covalent bonds govern intra-molecular organisation and are typically treated as fixed in modeling. In contrast, non-covalent, generally reversible interactions, including hydrogen bonds [50], salt bridges [51], van der Waals contacts [52], hydrophobic contacts [53], *π*–*π* stacking [54], and cation–*π* interactions [55], do not involve bond formation but arise from physicochemical compatibility between molecules. These interactions are predominantly inter-molecular and collectively determine protein–ligand binding, including affinity, specificity, and binding-site prediction. However, such local mechanistic details are typically absent from existing datasets. InteractBind addresses this limitation by explicitly representing such local interaction patterns through sequence token-level interaction maps.

#### S.1.2 Binding site and binding pocket

In structural biology, a <u>binding pocket</u> generally refers to a broader three-dimensional region on the protein surface that can accommodate a ligand, often characterized by geometric cavities or surface concavities [56]. In contrast, a <u>binding site</u> is more specific and denotes the subset of protein residues that directly participate in ligand recognition through non-covalent interactions [57]. While binding pockets describe a spatial region that may potentially host a ligand, binding sites capture the actual interacting components that determine molecular recognition and binding specificity. Although the two concepts are closely related and sometimes used interchangeably, they differ in granularity and interpretability. Binding pocket identification is often based on structural geometry and provides a coarse description of possible ligand-accessible regions, whereas binding site annotation reflects the underlying specific physicochemical interactions that stabilize binding [58]. In many sequence-based settings where high-resolution structural information is limited, binding pocket definitions can be ambiguous, making it difficult to evaluate whether a model has learned the true interaction determinants [19]. In this work, we therefore focus on binding sites at the residue level, defined by experimentally derived structural interaction annotations. This formulation provides a more precise and mechanistically grounded target for evaluation, enabling fine-grained assessment of whether models recover the actual residues involved in protein–ligand prediction tasks.

#### S.1.3 Biological background of ACE and L2HGDH

**ACE** is a central regulator of the renin–angiotensin system, catalyzing the conversion of angiotensin I to the vasoconstrictor angiotensin II and thereby controlling blood pressure, vascular tone, and cardiovascular homeostasis [59, 60]. Dysregulated ACE activity is a key driver of hypertension and cardiovascular disease, making ACE inhibition a cornerstone therapeutic strategy [38, 61]. As a result, extensive human ACE bioactivity data are available in public repositories such as PubChem [62] spanning a wide range of chemical scaffolds and potencies (Supplementary Fig. S3). This rich and well-characterized bioactivity landscape provides an ideal testbed for evaluating whether predictive models can reliably prioritize highly potent ligands based on inferred binding probabilities.

**L2HGDH** is a mitochondrial membrane–associated metabolic enzyme that catalyzes the oxidation of L-2-hydroxyglutarate (L-2-HG) to 2-oxoglutarate (2-OG); pathogenic mutations in *L2HGDH* impair this clearance and cause the neurometabolic disorder L-2-hydroxyglutaric aciduria (L-2-HGA) [63–65]. Genetic deficiency of L2HGDH in mice results in pathological accumulation of L-2-HG accompanied by leukoencephalopathy, neuroinflammation, and progressive neurodegeneration, consistent with the severe neurological phenotypes observed in human patients and motivating strategies aimed at restoring or enhancing L2HGDH activity [41]. In contrast, under acute cardiac ischemia and oxidative stress, L2HGDH loss–induced elevation of L-2-HG has been shown to preserve myocardial function by reprogramming glucose metabolism toward the pentose phosphate pathway and improving redox buffering, suggesting that transient and partial inhibition of L2HGDH may be beneficial in specific pathophysiological contexts [39]. Together, these findings establish L2HGDH as a context-dependent metabolic target for which both activation and inhibition are of potential therapeutic interest. In addition, L2HGDH is absent in the InteractBind training set and has no closely related homologs within it, enabling a stringent OOD evaluation of ExplainBind on an unseen target. Thus, we selected L2HGDH as an experimental (wet-lab) validation target and prioritized model-ranked compounds for experimental testing using in vitro assays of enzyme activity, enabling direct assessment of whether predicted binders act as functional inhibitors or activators of L2HGDH.

### S.2 Experimental setting

#### S.2.1 Curation of InteractBind dataset

##### Structure collection and sample construction

We construct **InteractBind**, a large-scale non-covalent interaction supervised protein–ligand binding database, by collecting protein–ligand complex structures from the Protein Data Bank (PDB) and processed them using PyMOL. Structures are filtered to retain only those that contain at least one organic ligand that binds to the protein. Each retained structure underwent a standardized preprocessing pipeline. All inorganic ligands and crystallographic water molecules are removed. Moreover, complexes containing metal ions in either the ligand or the protein are excluded. If a PDB entry contained multiple organic ligands, distance-based filtering is applied to extract individual ligand–protein complexes: for each ligand, only protein chains within 0.5 nm of the ligand are retained, while others are discarded. As a result, PDB files containing multiple ligands are split into multiple complexes, each consisting of a single ligand and its interacting protein chain(s).

For each complex, the input data were then separated into two files while preserving original atomic coordinates. Protein structures were converted to Structure-aware (SA)/FASTA format. Structure-aware (SA) protein representations were derived via *Foldseek* [43], providing an explicit local structural context [44]. Ligands were converted from PDB to SMILES and subsequently to SELFIES representations.

To obtain labeled binding data, a docking-based strategy was employed. Focused docking was performed for each ligand–protein pair, with the search box centred on the original ligand position and each axis set to the ligand size plus a 0.5 nm buffer. Pairs with predicted binding affinities lower (i.e., stronger) than *−*7.0 kcal*/*mol were labeled as positive samples. Negative samples were generated by randomly selecting ligands for each positive protein and performing global docking with the search box centered at the protein centroid and sized as the protein dimensions plus a 1.0 nm buffer. Pairs with top-ranked docking affinities higher (i.e., weaker) than *−*5.0 kcal*/*mol were labeled as negative samples. This procedure was repeated until each positive protein was associated with at least one valid negative ligand.

**Figure S1:**
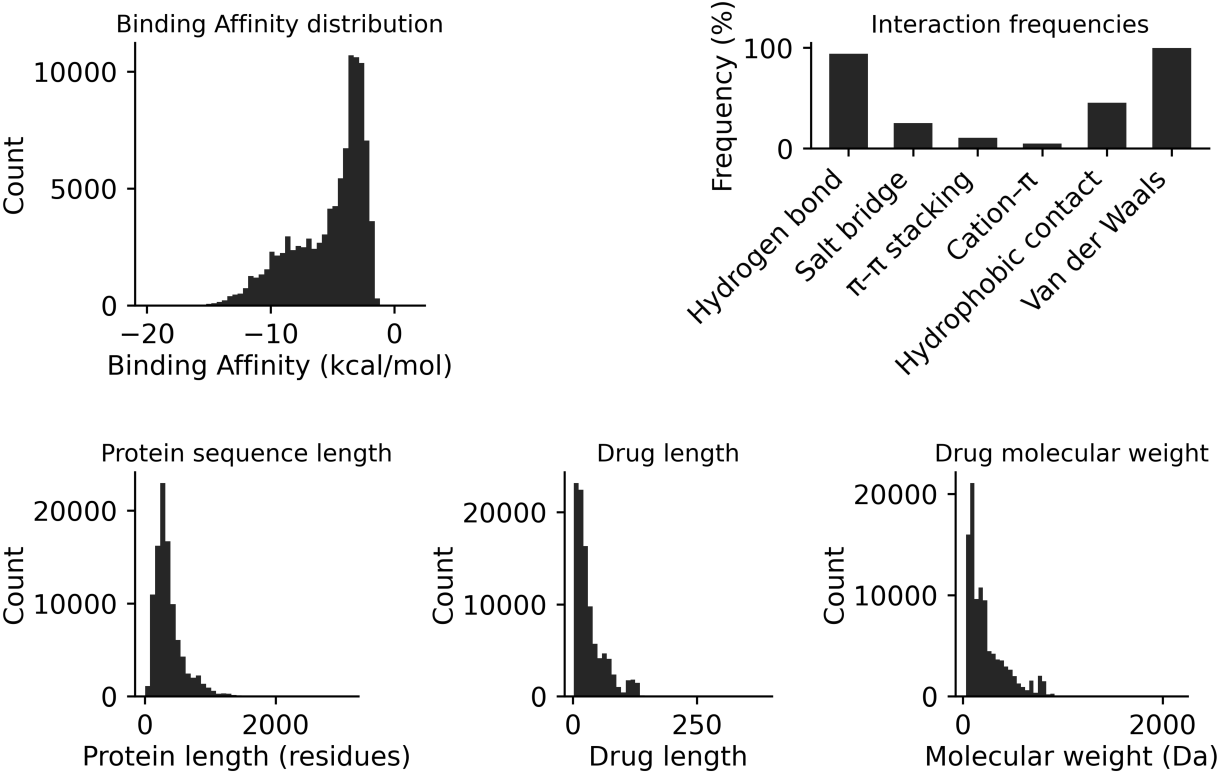
Distribution of features in **InteractBind**, including binding affinities, non-covalent interactions, protein sequence lengths, ligand lengths, and molecular weights.

##### Interaction annotation and strength modeling

Protein–ligand binding is governed by a diverse set of non-covalent interactions, including hydrogen bonds [50], salt bridges [51], van der Waals contacts [52], hydrophobic contacts [53], *π*–*π* stacking [54], and cation–*π* interactions [55]. These interactions jointly determine binding affinity, specificity, and structural complementarity, forming the physicochemical basis of molecular recognition [6]. Protein–ligand interactions for all positive samples in **InteractBind** are detected using rule-based criteria grounded in geometric and chemical constraints. Hydrogen bonds, salt bridges, *π*–*π* stacking, cation–*π* interactions, and hydrophobic contacts are identified using PLIP. van der Waals interactions, which are not captured by PLIP, are computed using GetContacts, and only attractive van der Waals interactions are retained. To quantify interaction strength, we adopt a distance-dependent piecewise linear decay function that maps interatomic or geometric distances to continuous values in the range [10^*−*6^, 1] within interaction-specific cutoff windows. Distance thresholds and geometric constraints for each non-covalent interaction are summarised in Table S1. Angular and geometric criteria, when applicable, are used as binary filters, while the distance-based decay controls the interaction strength.

Based on the detected interactions and their distance-dependent strengths, interaction-supervised attention maps are constructed at the token level. For each protein–ligand complex in **InteractBind**, an *m × n* matrix is generated, where *m* and *n* denote the numbers of ligand SELFIES tokens and protein FASTA tokens, respectively. Each matrix entry encodes the interaction strength between a ligand token and a protein residue. For each complex, six attention maps corresponding to individual non-covalent interactions are generated, together with an additional unified map obtained by summing all interaction-specific maps. All attention maps are stored using unique identifiers for efficient retrieval during training. After cleaning and preprocessing, **InteractBind** contains a total of **99**,**382 protein–ligand complexes**. These complexes form the basis for subsequent analyses and reflect substantial diversity in binding affinities, interaction patterns, and molecular properties, as shown in Fig. S1.

**Table S1:**
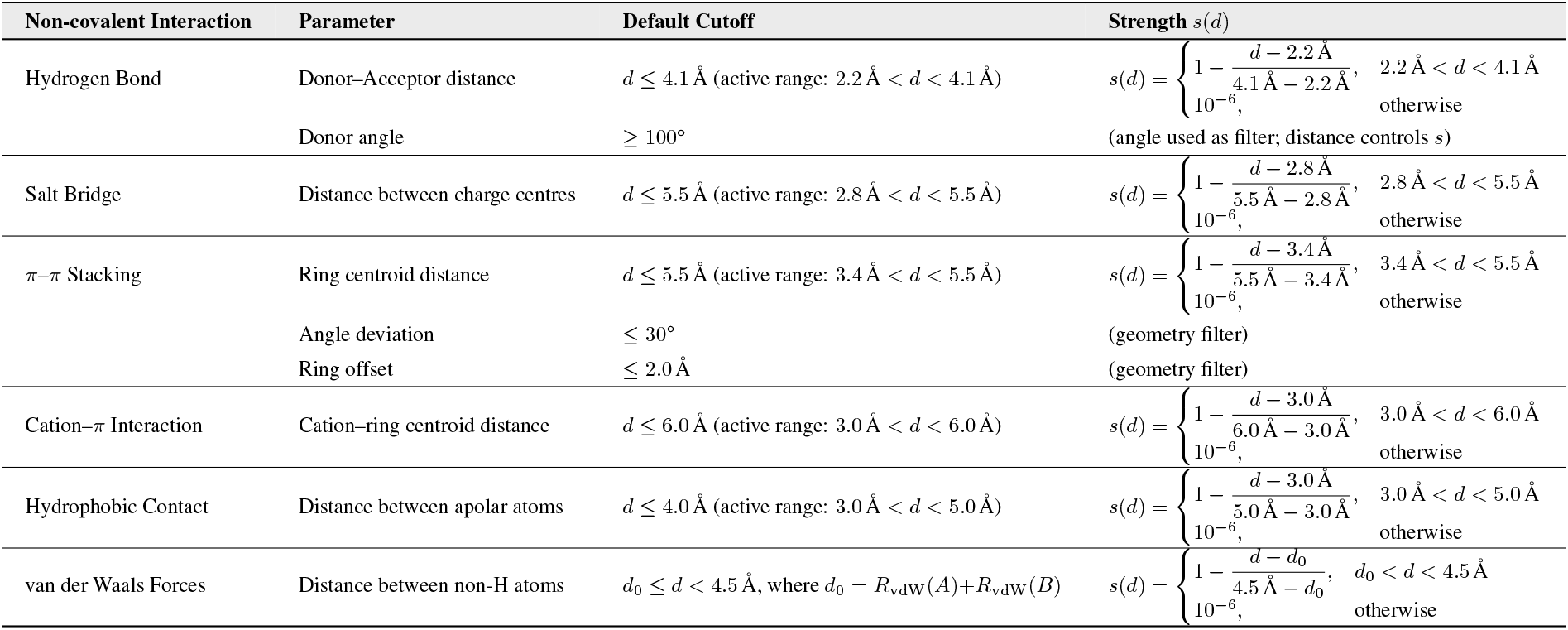
Cutoffs and strength definitions for key non-covalent interactions in InteractBind.

| Non-covalent Interaction | Parameter | Default Cutoff | Strength $s(d)$ |
| --- | --- | --- | --- |
| Hydrogen Bond | Donor–Acceptor distance | $d \leq 4.1 \text{ \AA}$ (active range: $2.2 \text{ \AA} < d < 4.1 \text{ \AA}$) | $s(d) = \begin{cases} 1 - \frac{d - 2.2 \text{ \AA}}{4.1 \text{ \AA} - 2.2 \text{ \AA}}, & 2.2 \text{ \AA} < d < 4.1 \text{ \AA} \\ 10^{-6}, & \text{otherwise} \end{cases}$ |
| | Donor angle | $\geq 100^\circ$ | (angle used as filter; distance controls $s$) |
| Salt Bridge | Distance between charge centres | $d \leq 5.5 \text{ \AA}$ (active range: $2.8 \text{ \AA} < d < 5.5 \text{ \AA}$) | $s(d) = \begin{cases} 1 - \frac{d - 2.8 \text{ \AA}}{5.5 \text{ \AA} - 2.8 \text{ \AA}}, & 2.8 \text{ \AA} < d < 5.5 \text{ \AA} \\ 10^{-6}, & \text{otherwise} \end{cases}$ |
| π–π Stacking | Ring centroid distance | $d \leq 5.5 \text{ \AA}$ (active range: $3.4 \text{ \AA} < d < 5.5 \text{ \AA}$) | $s(d) = \begin{cases} 1 - \frac{d - 3.4 \text{ \AA}}{5.5 \text{ \AA} - 3.4 \text{ \AA}}, & 3.4 \text{ \AA} < d < 5.5 \text{ \AA} \\ 10^{-6}, & \text{otherwise} \end{cases}$ |
| | Angle deviation | $\leq 30^\circ$ | (geometry filter) |
| | Ring offset | $\leq 2.0 \text{ \AA}$ | (geometry filter) |
| Cation–π Interaction | Cation–ring centroid distance | $d \leq 6.0 \text{ \AA}$ (active range: $3.0 \text{ \AA} < d < 6.0 \text{ \AA}$) | $s(d) = \begin{cases} 1 - \frac{d - 3.0 \text{ \AA}}{6.0 \text{ \AA} - 3.0 \text{ \AA}}, & 3.0 \text{ \AA} < d < 6.0 \text{ \AA} \\ 10^{-6}, & \text{otherwise} \end{cases}$ |
| Hydrophobic Contact | Distance between apolar atoms | $d \leq 4.0 \text{ \AA}$ (active range: $3.0 \text{ \AA} < d < 5.0 \text{ \AA}$) | $s(d) = \begin{cases} 1 - \frac{d - 3.0 \text{ \AA}}{5.0 \text{ \AA} - 3.0 \text{ \AA}}, & 3.0 \text{ \AA} < d < 5.0 \text{ \AA} \\ 10^{-6}, & \text{otherwise} \end{cases}$ |
| van der Waals Forces | Distance between non-H atoms | $d_0 \leq d < 4.5 \text{ \AA}$, where $d_0 = R_{\text{vdW}}(A) + R_{\text{vdW}}(B)$ | $s(d) = \begin{cases} 1 - \frac{d - d_0}{4.5 \text{ \AA} - d_0}, & d_0 < d < 4.5 \text{ \AA} \\ 10^{-6}, & \text{otherwise} \end{cases}$ |

#### S.2.2 Dataset splitting

We evaluate model performance on nine datasets derived from the InteractBind database under two complementary evaluation settings: in-distribution (ID) and out-of-distribution (OOD). These datasets are designed to assess both standard predictive accuracy and generalization under controlled distribution shifts. For ID evaluation, we construct the **InteractBind-ID** dataset. Unlike conventional PLB benchmarks that treat all protein–ligand complexes as positive samples, this dataset explicitly accounts for binding strength by stratifying interactions according to binding affinity. Protein–ligand pairs with high affinity are regarded as positive samples, whereas weakly binding pairs are treated as negative samples, yielding a more challenging and discriminative ID evaluation setting.

To assess generalisability systematically, we further construct eight OOD datasets by explicitly controlling sequence similarity between the training and test sets, as shown in Table S2. Each OOD dataset is divided into training, validation, and test splits with a ratio of 8:1:1. To examine the impact of protein sequence divergence, four protein-based OOD datasets are created: **InteractBind-P (25% OOD), InteractBind-P (28% OOD), InteractBind-P (31% OOD)**, and **InteractBind-P (33% OOD)**. In parallel, to evaluate ligand generalization, four ligand-based OOD datasets are constructed: **InteractBind-L (8% OOD), InteractBind-L (35% OOD), InteractBind-L (40% OOD)**, and **InteractBind-L (59% OOD)**. Compared with random in-distribution splits, these OOD settings provide a more realistic assessment of model generalization, reflecting practical drug discovery scenarios in which test proteins or molecules can differ substantially from those observed during training.

**Table S2:** Statistics of the InteractBind dataset, the in-distribution subset, and the protein- and ligand-similarity-controlled OOD datasets.

| Dataset | Setting | Split criterion | Pairs | Unique proteins | Unique ligands |
| --- | --- | --- | --- | --- | --- |
| InteractBind | Complete dataset | All curated pairs | 99,391 | 11,473 | 9,017 |
| InteractBind-ID | In-distribution | Affinity-based split | 84,825 | 10,942 | 6,911 |
| InteractBind-P (25% OOD) | Out-of-distribution | Protein similarity | 13,126 | 3,278 | 9,527 |
| InteractBind-P (28% OOD) | Out-of-distribution | Protein similarity | 13,126 | 2,966 | 8,957 |
| InteractBind-P (31% OOD) | Out-of-distribution | Protein similarity | 13,126 | 3,202 | 10,083 |
| InteractBind-P (33% OOD) | Out-of-distribution | Protein similarity | 13,126 | 4,807 | 11,084 |
| InteractBind-L (8% OOD) | Out-of-distribution | Ligand similarity | 7,994 | 3,999 | 6,132 |
| InteractBind-L (35% OOD) | Out-of-distribution | Ligand similarity | 7,994 | 3,102 | 2,114 |
| InteractBind-L (40% OOD) | Out-of-distribution | Ligand similarity | 7,994 | 2,926 | 1,577 |
| InteractBind-L (59% OOD) | Out-of-distribution | Ligand similarity | 7,994 | 2,186 | 715 |

#### S.2.3 Binding-site and interaction evaluation metrics: BRHR and IHR

We evaluate fine-grained protein–ligand predictions from two complementary perspectives: binding-site localization and non-covalent interaction prediction. For a given protein–ligand pair, let **P** *∈* ℝ^*m×n*^ denote the predicted interaction map, where *m* is the number of ligand tokens, *n* is the number of protein residue tokens, and *P*_*ij*_ is the predicted interaction score between ligand token *d*_*i*_ and protein residue token *p*_*j*_. We first rank all entries in **P** and select the Top-*K* predicted token pairs:

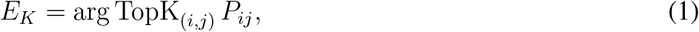

where *E*_*K*_ contains the index pairs of the *K* largest scores in **P**.

##### Binding-site localization

Binding-site localization evaluates whether the model identifies the correct protein-side binding region. Since this task only concerns protein residues, we project the predicted token pairs in *E*_*K*_ onto the protein axis:

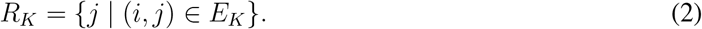

Let *T*_*R*_ denote the set of ground-truth binding residues. The sample-level Binding Residue Hit Rate (BRHR) is defined as

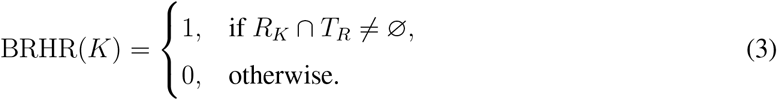

Thus, BRHR counts a prediction as correct if at least one predicted protein residue matches a ground-truth binding residue, regardless of the corresponding ligand token.

##### Non-covalent interaction prediction

Non-covalent interaction prediction evaluates whether the model recovers the annotated interaction at the token-pair level. Let *T*_*E*_ denote the set of ground-truth token pairs involved in non-covalent interactions. The sample-level Interaction Hit Rate (IHR) is defined as

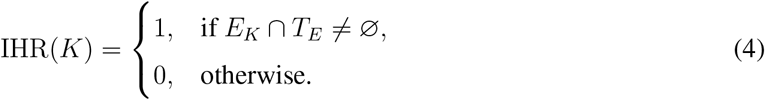

Unlike BRHR, IHR requires both sides of the predicted token pair to be correct, and therefore provides a stricter evaluation of non-covalent interaction prediction.

For both metrics, the dataset-level score is computed by averaging over all evaluated protein–ligand pairs:

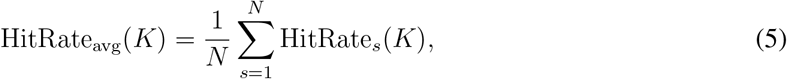

where HitRate denotes either BRHR or IHR, and *N* is the number of evaluated samples. The same protocol can be applied either to a unified interaction map or separately to interaction-specific maps for different non-covalent interaction types.

**Figure S2:**
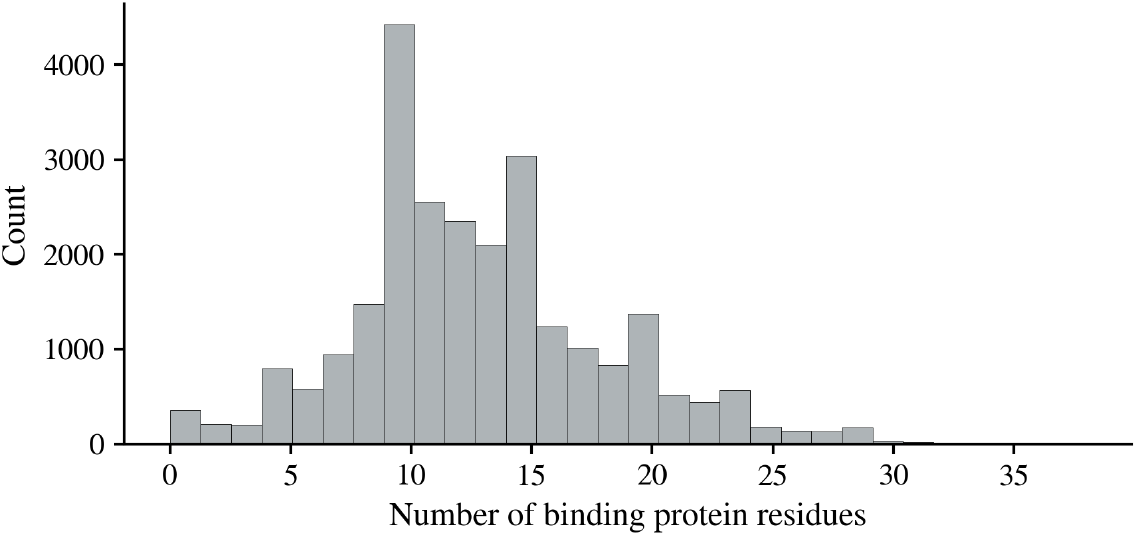
Distribution of ground-truth binding residues in the InteractBind dataset.

#### S.2.4 Baselines

We compare ExplainBind against the following seven representative baselines for the PLB task:

1. MolTrans [35], which tokenizes protein sequences and ligands into substructures, encodes them with transformer-based encoders, and applies an interaction module to model cross-modal binding patterns.
2. TransformerCPI [20], which represents proteins and compounds as sequences and employs transformer architectures to model their interactions for compound–protein interaction prediction.
3. HyperAttentionDTI [21], which introduces a hyper-attention mechanism to capture fine-grained interactions between protein residues and ligand atoms for drug–target interaction prediction.
4. PerceiverCPI [36], which adopts a Perceiver-based architecture to encode long protein and ligand sequences and model compound–protein interactions through latent cross-attention.
5. CAT-DTI [27], which uses cross-attention mechanisms to jointly model protein and ligand representations and improve interaction prediction.
6. DrugBAN [28], which encodes ligands with a graph neural network and proteins with a 1D CNN, and employs a bilinear attention network [66] to capture fine-grained residue–atom interactions before final prediction.
7. GraphBAN [37], which extends bilinear attention-based interaction modeling with graph neural representations for ligands and protein encoders for interaction prediction.

#### S.2.5 Hyperparameters

ExplainBind is implemented in Python 3.8 and the PyTorch framework (1.12.1)^1^. The computing device we use is the NVIDIA GeForce RTX A6000. Table S3 shows the parameters of the ExplainBind model and Table S4 lists the notations used in this paper with descriptions.

**Table S3:**
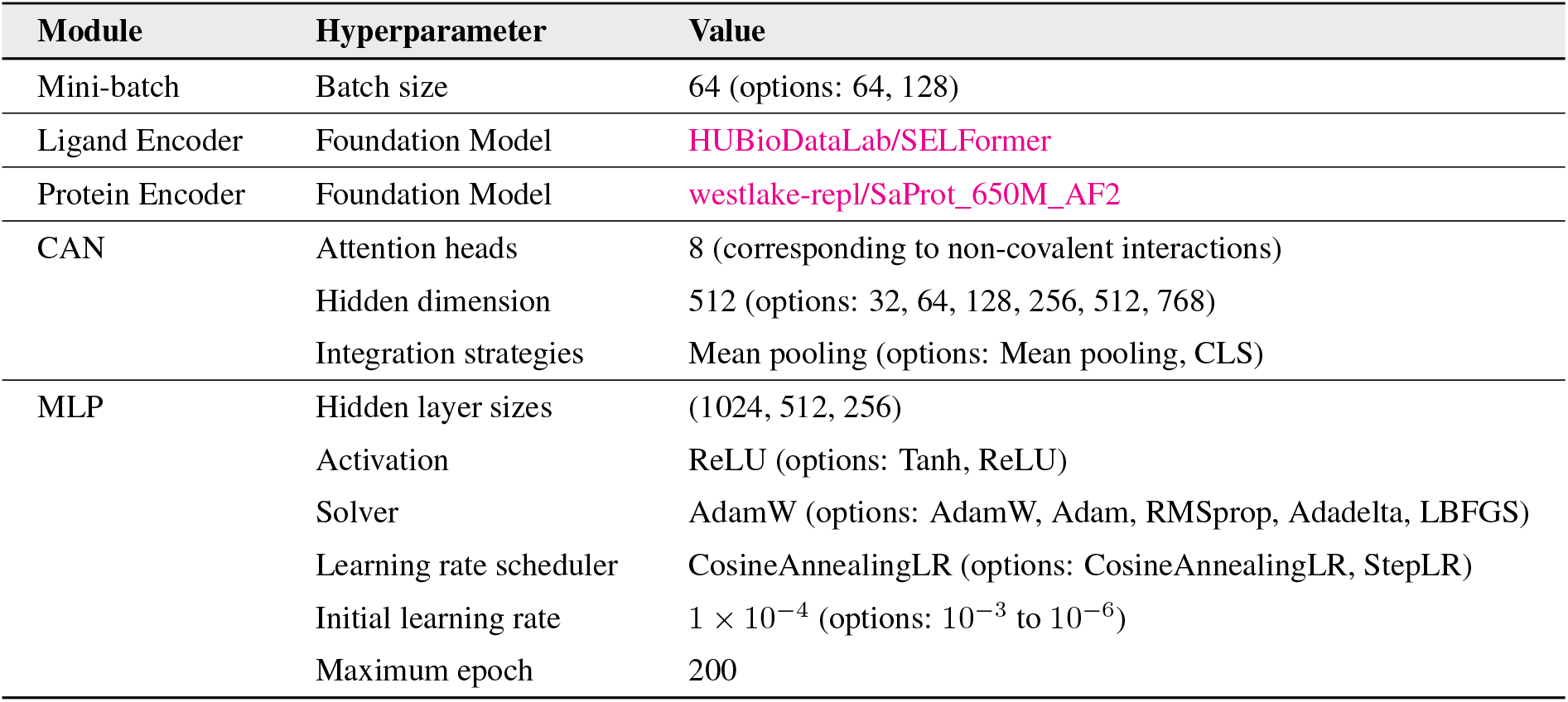
Configuration parameters of ExplainBind.

#### S.2.6 Curation of ACE dataset

ACE bioactivity data are curated from PubChem [62] in July 2024. Only human ACE bioassays reporting IC_50_ values are retained, while non-human assays and alternative bioactivity readouts are excluded. For compounds associated with multiple IC_50_ measurements, values with small relative variation are averaged, whereas compounds exhibiting large inter-assay discrepancies are removed to ensure data reliability and consistency. The corresponding human ACE amino acid sequence is retrieved from the UniProt database (P12821).

**Table S4:** Notations and descriptions used in ExplainBind.

| Notation | Description |
| --- | --- |
| $\mathbf{D} \in \mathbb{R}^{m \times h}$ | Ligand token embeddings (length $m$, hidden size $h$) |
| $\mathbf{P} \in \mathbb{R}^{n \times h}$ | Protein token embeddings (length $n$, hidden size $h$) |
| $\mathbf{Q}_d, \mathbf{K}_d, \mathbf{V}_d$ | Query, key, and value matrices for ligand tokens |
| $\mathbf{Q}_p, \mathbf{K}_p, \mathbf{V}_p$ | Query, key, and value matrices for protein tokens |
| $\mathbf{W}_q^d, \mathbf{W}_k^d, \mathbf{W}_v^d$ | Projection weights for ligand Q/K/V |
| $\mathbf{W}_q^p, \mathbf{W}_k^p, \mathbf{W}_v^p$ | Projection weights for protein Q/K/V |
| $\hat{\mathbf{A}}_{dp}^{(t)} \in \mathbb{R}^{m \times n}$ | Predicted ligand→protein attention map for head $t$ |
| $\hat{\mathbf{A}}_{pd}^{(t)} \in \mathbb{R}^{n \times m}$ | Predicted protein→ligand attention map for head $t$ |
| $\mathbf{A}^{(t)} \in \mathbb{R}^{m \times n}$ | Ground-truth ligand→protein interaction map for head $t$ |
| $\mathbf{A}_{\text{raw}}$ | Unnormalised ground-truth interaction intensity used to construct $\{\mathbf{A}^{(t)}\}$ |
| $\mathbf{P}_{\text{sa}}^{(t)}, \mathbf{D}_{\text{sa}}^{(t)}$ | Self-attended protein/ligand features for head $t$ |
| $\mathbf{P}^{*(t)}, \mathbf{D}^{*(t)}$ | Head-specific fused representations (cross + self fusion) |
| $\mathbf{P}^*, \mathbf{D}^*$ | Final fused sequences after concatenation across all heads |
| $\bar{\mathbf{D}}^*, \bar{\mathbf{P}}^*$ | Mean-pooled ligand/protein fused embeddings |
| $\mathbf{F} = [\bar{\mathbf{D}}^*, \bar{\mathbf{P}}^*]$ | Joint protein–ligand representation for classification |
| $p \in [0, 1]$ | Predicted binding probability |
| $\mathcal{L}_{\text{cls}}$ | Binary cross-entropy classification loss |
| $\mathcal{L}_{\text{att}}$ | Masked KL-divergence attention alignment loss |
| $\mathcal{L}$ | Total loss $(1 - \lambda)\mathcal{L}_{\text{cls}} + \lambda\mathcal{L}_{\text{att}}$ |
| $\mathcal{T}$ | Set of attention heads ($\vert \mathcal{T}\vert = 8$: six type-specific and two overall) |
| $m, n$ | Sequence lengths of ligand and protein |
| $h$ | Hidden dimension size |
| $\lambda$ | Weight balancing classification and attention losses |

**Figure S3:**
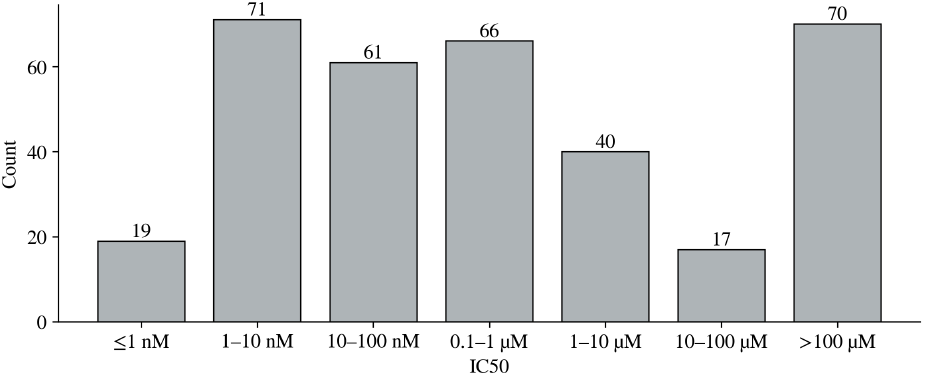
Distribution of ACE compound potency. Histogram of IC_50_ values for the curated human ACE dataset, grouped into logarithmic activity bins ranging from sub-nanomolar to >100 *µ*M.

The final curated dataset comprises 344 compounds with experimentally measured IC_50_ values spanning from *≤* 1 nM to *>* 100 *µ*M, thereby covering more than five orders of magnitude in inhibitory potency. As shown in Fig. S3, the activity distribution is highly heterogeneous across potency regimes. Highly potent inhibitors (IC_50_ *≤* 1 nM) are relatively rare, whereas the majority of compounds fall within the 1–10 nM, 10–100 nM, and sub-micromolar ranges. In contrast, a substantial fraction of weakly active or inactive compounds (IC_50_ *>* 100 *µ*M) is also present.

This long-tailed and imbalanced potency distribution mirrors realistic medicinal chemistry screening data and provides a stringent testbed for evaluating the ability of ExplainBind to prioritise highly potent ligands from a large background of weaker binders.

#### S.2.7 L2HGDH bioassay setup

The expression construct for the *Drosophila melanogaster* orthologue of L2HGDH (dmL2HGDH) with N-terminal histidine and SUMO tags is provided by Jianping Ding [65]. The protein is expressed and purified as previously described [65] using B-PER Complete (Thermo) with HALT Protease Inhibitor Cocktail (Thermo) for bacterial cell lysis. Purity is confirmed by UV-visible spectroscopy and mass spectrometry. For the activity assay, 10 *µ*L of 40 nM dmL2HGDH in assay buffer (20 mM NaH_2_PO_4_, pH 7.4, 0.05% CHAPS, and 1% DMSO) is incubated with test compounds at 20 *µ*M for 30 min at ambient temperature. The reaction is initiated by the addition of 10 *µ*L substrate solution containing 120 *µ*M L-2-hydroxyglutarate and 200 *µ*M resazurin in assay buffer. Following 90 min incubation, fluorescence intensity is measured using a PHERAstar FSX plate reader (BMG LABTECH) with excitation at 540 nm and emission at 590 nm. With this assay, the top 200 predicted molecules are tested in duplicate. Compound activity is calculated as the percentage relative to the DMSO vehicle control.

Relative activity is defined as the percent change in enzyme activity in the presence of compound relative to the vehicle control:

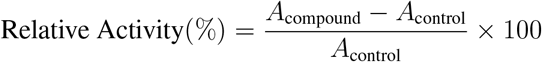

where *A*_compound_ is the measured enzyme activity in the presence of compound, and *A*_control_ is the activity measured with vehicle control. Positive values indicate activation, whereas negative values indicate inhibition. Absolute relative activity (|Relative Activity|) is used to quantify the magnitude of functional modulation regardless of direction. The amino acid sequence used for model evaluation is derived from the crystal structure of dmL2HGDH (PDB ID: 8W75).

#### S.2.8 Docking for L2HGDH binders

Molecular docking is performed to evaluate the binding feasibility and plausible binding modes of candidate ligands to human L2HGDH using AutoDock Vina [7]. The enzyme conformation is derived from the L2HGDH crystal structure (PDB: 8W78, chain A) and prepared in PDBQT format with the flavin adenine dinucleotide (**FAD**) cofactor retained to preserve the native catalytic environment. Ligands are converted to PDBQT format and docked individually in a batch workflow using a blind-docking strategy. To ensure consistent docking across all ligands, a single docking search space is defined directly from the receptor structure. All ATOM and HETATM coordinates in the receptor PDBQT file are used to determine the axis-aligned bounding box of the receptor. The docking box center is set to (8.43, *−*34.30, *−*18.29) Å, and the box dimensions are defined as 58.52 *×* 69.63 *×* 67.78 Åalong the *x, y*, and *z* axes, respectively. These dimensions correspond to the full receptor coordinate span expanded by a fixed padding of 5 Å, ensuring coverage beyond a tight protein envelope and accommodating surface-accessible binding sites. For visualization, the protein and docking search space are rendered using py3Dmol [67]. The protein is displayed as a light-gray schematic representation; hetero atoms including the retained FAD cofactor are shown as sticks, and the docking box is overlaid as a gray wireframe cuboid. The view is automatically centered and zoomed to the structure to enable rapid inspection of docking box placement and coverage prior to large-scale docking.

### S.3 Additional results

#### S.3.1 Non-covalent interaction prediction

We report the quantitative results for the fine-grained non-covalent interaction prediction setting defined in Section S.2.3. Unlike binding-site localisation, which evaluates whether predicted ligand–protein token pairs recover protein-side binding residues after projection onto the protein axis, this setting uses a stricter token-pair criterion. A prediction is counted as correct only when the predicted ligand-token–protein-residue pair matches a ground-truth interaction pair for the corresponding annotated interaction type.

**Table S5:** Interaction-specific IHR@*K* and Recall@*K* for non-covalent interaction prediction. The **highest** and <u>second-highest</u> values in each column are highlighted.

| Interaction type | IHR@1 | IHR@5 | IHR@10 | IHR@15 | Recall@5 | Recall@10 | Recall@15 |
| --- | --- | --- | --- | --- | --- | --- | --- |
| van der Waals forces | 1.9±0.3 | 10.3±0.9 | 19.5±1.4 | 26.7±1.6 | 0.9±0.2 | 2.1±0.3 | 3.1±0.4 |
| Hydrogen bonding | 1.6±0.2 | 5.4±0.7 | 10.6±1.0 | 15.5±1.2 | 1.3±0.2 | 2.8±0.4 | 4.4±0.5 |
| Salt bridges | <u>3.9±0.5</u> | <u>16.5±1.3</u> | 27.0±1.5 | 36.2±1.8 | <u>3.9±0.5</u> | <u>7.8±0.8</u> | <u>11.8±1.0</u> |
| π-π stacking | **7.3±0.6** | **29.8±1.8** | **44.2±2.0** | **52.5±2.3** | **5.7±0.6** | **11.2±1.0** | **16.3±1.3** |
| Cation-π interactions | 2.5±0.4 | 14.7±1.2 | <u>29.4±1.7</u> | <u>37.8±1.9</u> | 2.8±0.4 | 6.7±0.7 | 10.3±0.9 |
| Hydrophobic contacts | 1.7±0.3 | 6.7±0.7 | 12.9±1.1 | 17.2±1.3 | 1.4±0.3 | 3.2±0.4 | 4.5±0.5 |

Table S5 reports the interaction-specific IHR@*K* and Recall@*K* results. Across all interaction categories, performance consistently improves as *K* increases, indicating that correct ligand–protein interaction pairs are progressively recovered among the highest-ranked predictions. Among the six interaction types, *π*–*π* stacking achieves the strongest performance across all metrics, reaching an IHR@15 of 52.5% and Recall@15 of 16.3%. Salt bridges obtain the second-best results for IHR@1, IHR@5 and all Recall metrics, while cation–*π* interactions become the second-best category at larger hit-rate thresholds, reaching an IHR@10 of 29.4% and IHR@15 of 37.8%. Van der Waals forces show moderate recovery at larger *K*, whereas hydrogen bonding and hydrophobic contacts remain more challenging. This pattern suggests that the model more effectively identifies interaction types with distinctive physicochemical signatures, while sparse or geometrically constrained contacts are harder to recover at the token-pair level. Overall, these results demonstrate that the model can recover fine-grained, interaction-specific ligand–protein contact patterns beyond coarse binding-site localisation.

#### S.3.2 Case study details

A detailed analysis is presented below:

- **Homo sapiens CDK2–staurosporine** (Fig. 2d). The model correctly highlighted three non-covalent interactions—hydrogen bonds, hydrophobic interactions, and van der Waals interactions—that collectively anchor staurosporine within the CDK2 binding cleft. Key hydrogen bond donors or acceptors included GLU81, LEU83, ASP86, and GLN131, engaging the ligand’s indolocarbazole backbone in close agreement with crystallographic data. Hydrophobic interactions are concentrated around ALA31, LYS33, PHE80, and ASP145, forming a compact apolar cage that stabilizes the planar ring system. In addition, ILE10, GLY11, VAL64, PHE82, LEU83, HIS84, GLN85, and LEU134 contributed to van der Waals interactions along the aromatic scaffold, further reinforcing ligand binding within the pocket.
- **Sus scrofa elastase–4E4** (Fig. 2e). The ground-truth interaction map comprises four non-covalent interactions—hydrogen bonds, hydrophobic interactions, ***π*–*π*** stacking, and van der Waals interactions—all of which are accurately recovered by ExplainBind. The model identified hydrogen bonds between the inhibitor’s carbonyl and amide groups and residues GLN185, SER188, and HIS45 within the catalytic triad region. Hydrophobic contacts clustered around VAL88, VAL209, and PHE208 provide the principal non-polar anchoring surface. Notably, HIS45 is also enriched in *π*–*π* stacking, highlighting dual aromatic–electrostatic stabilization within the substrate-recognition pocket. In addition, HIS45, CYS184, and GLN185 contributed to van der Waals interactions, further reinforcing local packing between the ligand and the catalytic groove.
- **Staphylococcus aureus DHFR–trimethoprim** (Fig. 2f). The model successfully reproduced the hydrogen bond network between the ligand’s amino and methoxy groups and the enzyme’s active-site residues LEU5, ASP27, and PHE92. Hydrophobic interactions are correctly localized to ILE50, LEU20, and PHE92, which surround the *p*-aminobenzyl ring of trimethoprim and stabilize its orientation within the pocket. Additionally, LEU5, VAL6, ALA7, LEU20, SER49, and PHE92 are predicted to contribute to van der Waals interactions, collectively forming a mixed polar–apolar microenvironment characteristic of the DHFR binding site.

#### S.3.3 Prioritization of high-activity ligands with wet-lab validation on L2HGDH

Table S6 presents a Top-*K* analysis of model predictions for ligand prioritization in the L2HGDH case study. Ligands are ranked by predicted binding probability, and we observe a clear enrichment of experimentally active molecules among the highest-ranked candidates. In particular, the mean |Relative Activity is substantially higher for small Top-*K* subsets and decreases monotonically as *k* increases, while the mean predicted probability remains close to unity for the top-ranked ligands. Consistently, molecules with |Activity| *≥* 25 are predominantly concentrated within the high-confidence region, supporting the use of model confidence for guiding candidate selection in subsequent wet-lab validation.

**Table S6:**
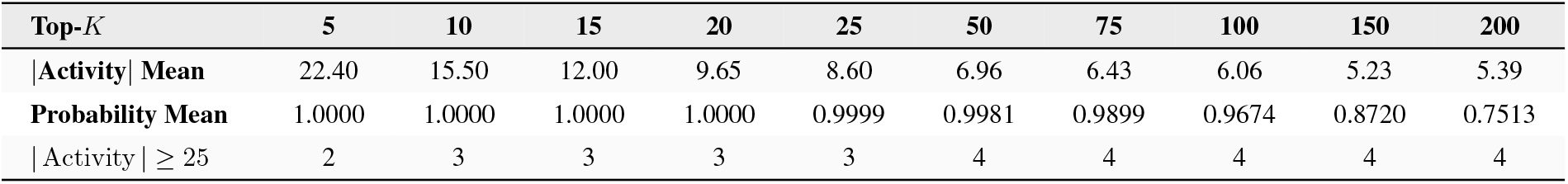
Top-*K* analysis of model predictions. We report the mean |Relative Activity, mean predicted probability, and the number of samples with |Activity| *≥* 25 for each Top-*K*.

#### S.3.4 Efficiency analysis

Efficiency is a critical factor when scaling computational models to large drug discovery datasets, and it is strongly influenced by how molecular representations are managed. In ExplainBind, the ligand and protein encoders are frozen, so their embeddings remain unchanged during training. This enables a pre-encoding strategy, where all representations are computed once in advance and stored for reuse. In contrast, baselines such as DrugBAN and GraphBAN continuously update embeddings during training and therefore cannot benefit from this approach. As illustrated in Fig. S4, on the InteractBind-ID dataset the training time of ExplainBind without pre-encoding (about 220 minutes) is comparable to that of the bilinear baselines (DrugBAN *∼*230 minutes; GraphBAN *∼*250 minutes). With pre-encoding, however, the training time of ExplainBind is reduced to approximately 85 minutes. The bar chart clearly demonstrates this substantial gain in efficiency, which arises from avoiding repeated feature extraction. A limitation of pre-encoding is the additional memory required to store pre-computed embeddings, which may become a consideration when scaling to very large datasets.

**Figure S4:**
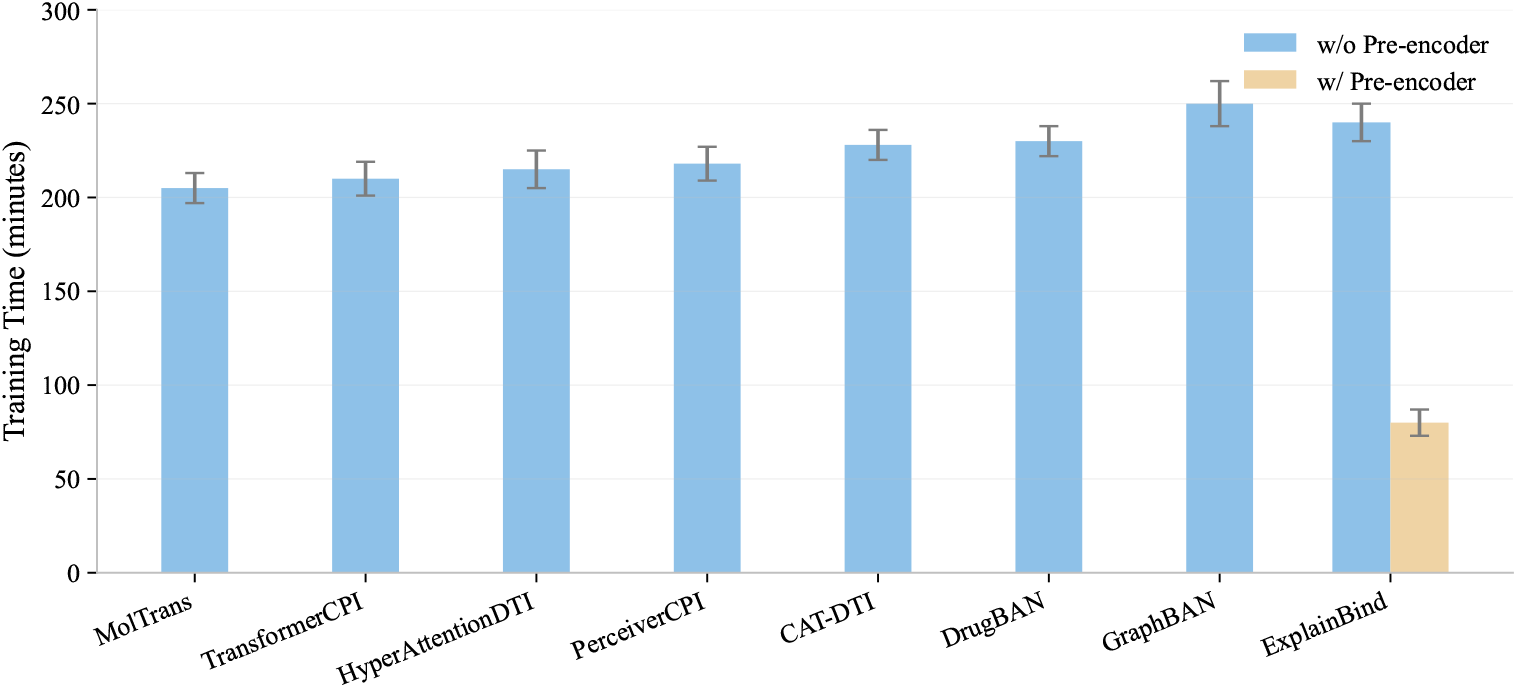
Comparison of training time on the InteractBind-ID dataset.

#### S.3.5 Time complexity analysis

The feature dimensions of the representations generated by different foundation model encoders are fixed, but the dimensionality may differ across encoders. To enable fusion of protein and ligand representations, we use linear layers to project them into a common hidden dimension (*h*) of 512.

For the token-level interaction in the PLB task, the time complexity is dominated by the attention mechanism. With multi-head attention (*H* = 8), the computation of queries, keys, and values, as well as the softmax attention weights, requires

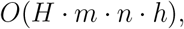

where *m* and *n* are the sequence lengths of ligand and protein tokens, respectively, and *h* is the hidden dimension. Since each head attends across all pairs of residue–level tokens, the *m · n* term (from the pairwise attention score computation) is the key factor. Thus, the overall time complexity of the supervised attention module is

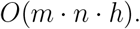

**Table S7:** Time complexity and parameter count of the supervised attention network (CAN).

| Module | Complexity (O) | Parameters |
| --- | --- | --- |
| CAN | $O(m \cdot n \cdot h)$ | 1.57M |

In summary, the supervised attention network scales linearly with the hidden dimension and quadratically with the token lengths of protein and ligand sequences. While this introduces higher parameter counts compared to simpler bilinear modules, the explicit attention supervision enables mechanistically explainable residue–level interaction maps without prohibitive computational overhead.

#### S.3.6 Evaluation of different encoders

##### Descriptions

- **ProteinBERT** [68]: Pre-trained on large-scale amino acid sequences to capture evolutionary and contextual dependencies within protein families.
- **ESM-2** [45]: An evolutionary-scale protein language model trained on UniRef50, offering deep contextual representations from protein sequence space.
- **SaProt** [44]: Incorporates 3D structural priors by transforming protein structures into structure-aware sequences via FoldSeek.
- **ChemBERTa-2** [49]: A transformer-based molecular encoder trained on large SMILES corpora, providing chemically informed embeddings.
- **MoLFormer** [48]: A multi-task transformer model pre-trained on both molecular graphs and SMILES strings, enabling enhanced molecular generalization.
- **SELFormer** [46]: Utilises SELFIES as a chemically valid representation, improving robustness and representation stability in molecular encoding.

**Table S8:** Summary of foundation models used for encoding proteins and ligands. Protein encoders process either amino acid or structure-aware sequences, while ligand encoders accept SMILES or SELFIES representations.

| Foundation Models | Model Name | HuggingFace Link | Input Type |
| --- | --- | --- | --- |
| Protein | ProteinBERT | Rostlab/prot_bert | Amino acid sequence |
|  | ESM-2 | facebook/esm2_t33_650M_UR50D | Amino acid sequence |
|  | SaProt | westlake-repl/SaProt_650M_AF2 | Structure-aware sequence |
| Ligand | ChemBERTa-2 | seyonec/ChemBERTa-zinc-base-v1 | SMILES |
|  | MoLFormer | ibm-research/MoLFormer-XL-both-10pct | SMILES |
|  | SELFormer | HUBioDataLab/SELFormer | SELFIES |

Our framework supports flexible substitution of these encoders with emerging state-of-the-art foundation models, ensuring continual adaptability and performance improvement as the field evolves.

**Table S9:** Performance comparison of different protein and ligand encoder combinations on the PDB (33% OOD) dataset. Ligand inputs are SMILES or SELFIES, while protein inputs are amino acid sequences or structure-aware representations. We highlight the **best** and <u>second best</u> results.

| Ligand Input | Protein Input | Ligand Encoder | Protein Encoder | Accuracy | AUROC | AUPRC |
| --- | --- | --- | --- | --- | --- | --- |
| SMILES | Amino acid sequence | ChemBERTa-2 | ProteinBERT | 0.796±0.007 | 0.902±0.004 | 0.894±0.005 |
| SMILES | Amino acid sequence | ChemBERTa-2 | ESM-2 | 0.794±0.006 | 0.900±0.003 | 0.896±0.004 |
| SMILES | Structure-aware sequence | ChemBERTa-2 | SaProt | 0.807±0.005 | 0.927±0.003 | 0.899±0.003 |
| SMILES | Amino acid sequence | MoLFormer | ProteinBERT | 0.787±0.006 | 0.914±0.005 | 0.902±0.003 |
| SMILES | Amino acid sequence | MoLFormer | ESM-2 | 0.792±0.005 | 0.917±0.004 | 0.901±0.006 |
| SMILES | Structure-aware sequence | MoLFormer | SaProt | <u>0.811±0.005</u> | <u>0.945±0.003</u> | <u>0.927±0.008</u> |
| SELFIES | Amino acid sequence | SELFormer | ProteinBERT | 0.798±0.003 | 0.930±0.004 | 0.912±0.006 |
| SELFIES | Amino acid sequence | SELFormer | ESM-2 | 0.805±0.006 | 0.940±0.004 | 0.926±0.005 |
| SELFIES | Structure-aware sequence | SELFormer | SaProt | **0.813±0.004** | **0.947±0.003** | **0.929±0.002** |

Accurate token-level fusion is critical for the representational quality of both protein and ligand encoders. As summarized in Table S9, the best overall performance is achieved by combining SELFIES ligands encoded by SELFormer with SA protein sequences encoded by SaProt. This combination achieves the highest scores across all metrics, reflecting the advantages of self-consistent molecular representations on both sides. It is worth noting that SA sequences require extracting structure-aware features from protein structures using FoldSeek [43]. For proteins without experimentally determined structures, we also provide a model trained on the SELFIES & amino-acid sequence combination, which allows users to perform inference using only sequence-level input. This ensures compatibility with sequence-only datasets while maintaining strong performance. In practice, users can choose the appropriate model according to the available molecular inputs. For ligands, both **SMILES** and **SELFIES** are supported, while for proteins, either **structure-aware** or **amino acid** sequences can be used. As indicated in Table S9, the performance degradation from structure-aware to sequence-only setting is limited, confirming the robustness and generalisability of ExplainBind’s foundation encoders across different molecular input formats.

#### S.3.7 Performance on public datasets of protein-ligand binding prediction

To assess further the robustness of ExplainBind, we additionally conducted experiments on three widely used public datasets: **BindingDB** [69], **BioSNAP** [70], and **Human** [20, 71]. In the ID setting, the model is trained solely with binary interaction labels, without the guidance of residue–level attention maps. As shown in Fig. S5, ExplainBind consistently outperforms a wide range of 8 baselines, indicating that it maintains strong predictive capability even without explicit structural supervision.

We further evaluated ExplainBind under OOD scenarios, where training and test sets differ substantially in protein and ligand distributions. The results demonstrate that ExplainBind achieves the best performance in two OOD settings and ranks second in the remaining case, presented in Fig. S6. These findings highlight that ExplainBind generalises well across heterogeneous benchmarks and remains competitive under challenging distribution shifts.

**Figure S5:**
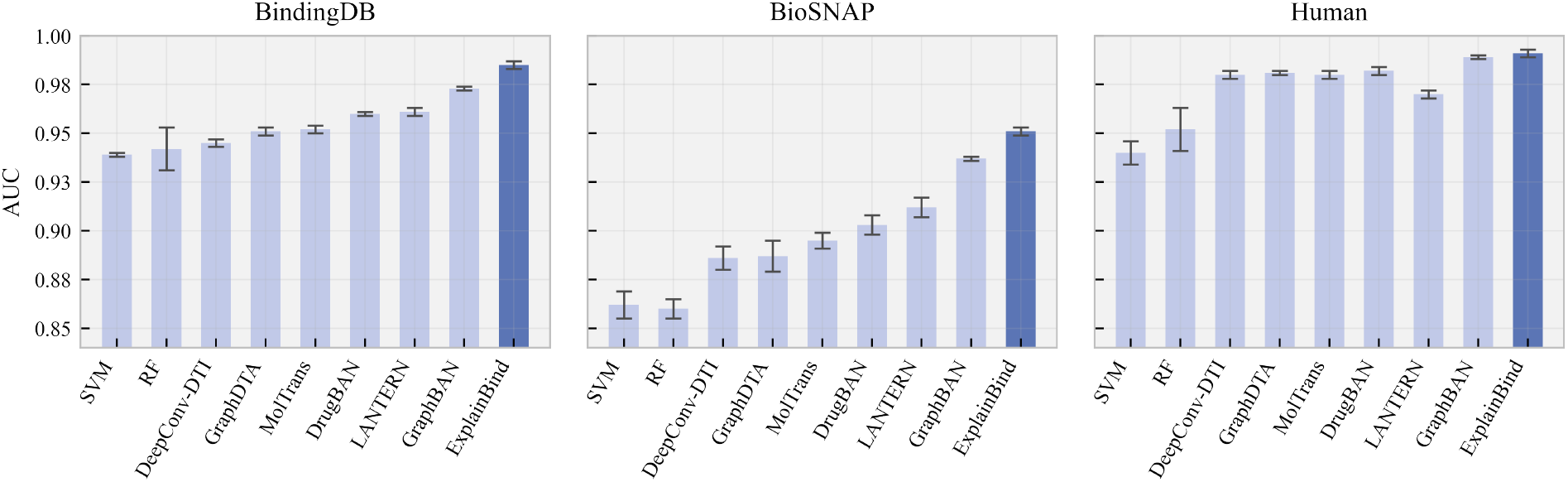
In-distribution performance comparison on three publicly available datasets without attention map supervision.

#### S.3.8 Ablation study

To evaluate the contribution of individual components in ExplainBind, we conducted a comprehensive ablation analysis as summarized in Table S10. All variants share the same configuration as the final ExplainBind model. Replacing the pooling-based fusion with a [CLS] aggregation strategy results in a notable drop in performance (AUROC 0.940 *→* 0.927; Accuracy 0.805 *→* 0.784), confirming that mean pooling offers a more stable integration of token-level evidence. When replacing FASTA sequences with SA protein representations, the model achieves the best overall results (AUROC 0.947; Accuracy 0.817), suggesting that structural context can further strengthen the learned protein–ligand interactions. However, the SA variant requires structural features extracted through *Foldseek*, which depend on the availability of experimentally resolved or high-confidence predicted protein structures. In practice, many proteins are represented only by their amino acid sequences, making the FASTA-based ExplainBind a more universally applicable model. Overall, these findings demonstrate that pooling aggregation contributes to ExplainBind’s robust and explainable performance. In Supplementary S.3.6, we further provide detailed comparisons of alternative protein–ligand encoder configurations.

**Figure S6:**
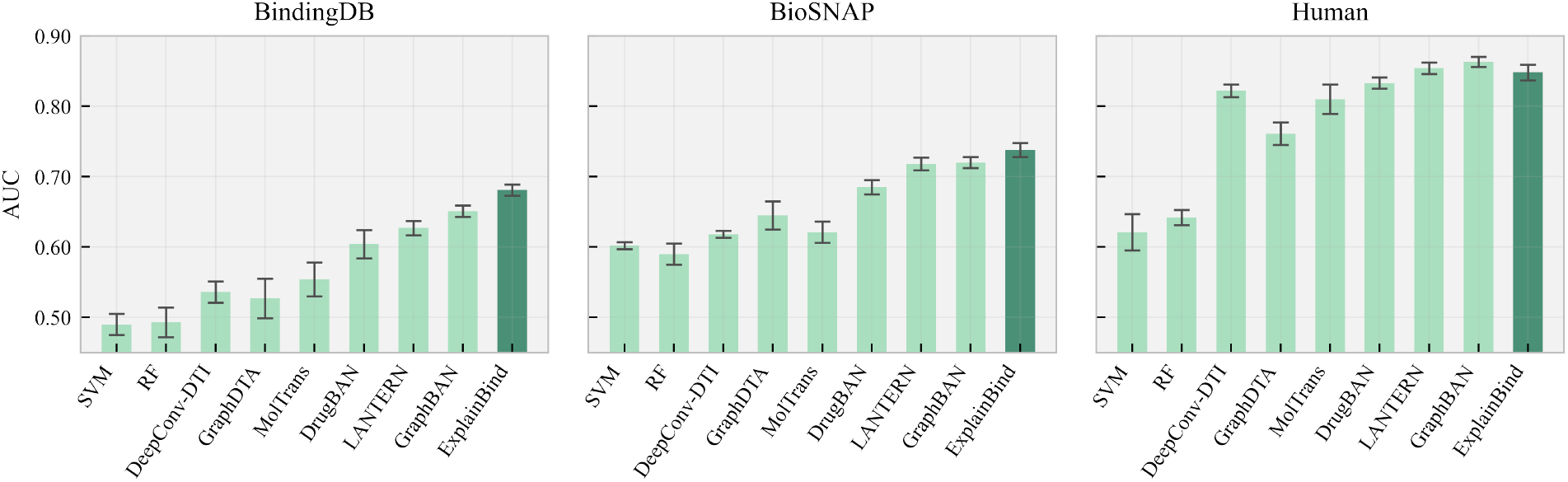
Out-of-distribution performance comparison on three publicly available datasets without attention map supervision.

**Table S10:** Ablation results on the PDB (33% OOD) dataset. All results are based on the same ExplainBind configuration; each row removes or replaces one component for comparison.

| Model / Setting | AUROC | AUPRC | Accuracy | Description |
| --- | --- | --- | --- | --- |
| w/o Attn Sup. | 0.943 | 0.930 | 0.812 | removing attention-map supervision |
| w/ Attn Sup. | 0.940 | 0.926 | 0.805 | with attention-map supervision |
| CLS Aggregation | 0.927 | 0.913 | 0.784 | using [CLS] aggregation (↓ vs ExplainBind) |
| Pooling Aggregation | 0.940 | 0.926 | 0.805 | using mean pooling for fusion |
| ExplainBind (SA) | 0.947 | 0.933 | 0.817 | variant using structure-aware protein representation (↑ vs ExplainBind) |
| ExplainBind | 0.940 | 0.926 | 0.805 | using raw FASTA sequence input |
| KL ratio = 0.1 | 0.937 | 0.920 | 0.794 | varying KL loss ratio for attention alignment |
| KL ratio = 0.2 | 0.936 | 0.923 | 0.797 |  |
| KL ratio = 0.3 | **0.940** | **0.926** | **0.805** |  |
| KL ratio = 0.4 | 0.939 | 0.923 | 0.781 |  |
| KL ratio = 0.5 | 0.935 | 0.921 | 0.779 |  |

#### S.3.9 OOD performance comparison

Table S11 reports the exact numerical results underlying the OOD performance line plots presented in the main text. Specifically, the table summarises the OOD performance of ExplainBind and baseline methods on the InteractBind-P datasets under varying OOD ratios (25–33%), evaluated using multiple metrics: AUROC, AUPRC, Accuracy, F1 score, Sensitivity, Specificity, and MCC [72].

**Table S11:** OOD performance comparison of ExplainBind and baselines. The best and <u>second best</u> results are highlighted. InteractBind-P (25–33% OOD) denotes the out-of-distribution test sets.

| Model | AUROC | AUPRC | Accuracy | F1 | Sensitivity | Specificity | MCC |
| --- | --- | --- | --- | --- | --- | --- | --- |
| **InteractBind-P (33% OOD)** — Out-of-distribution datasets (decreasing dissimilarity) |  |  |  |  |  |  |  |
| MolTrans | 0.795±0.015 | 0.765±0.016 | 0.678±0.012 | 0.668±0.013 | 0.660±0.015 | 0.682±0.012 | 0.230±0.013 |
| TransformerCPI | 0.822±0.014 | 0.784±0.015 | 0.692±0.012 | 0.681±0.013 | 0.675±0.014 | 0.696±0.012 | 0.241±0.012 |
| HyperAttentionDTI | 0.823±0.013 | 0.798±0.014 | 0.704±0.011 | 0.694±0.012 | 0.688±0.013 | 0.708±0.011 | 0.251±0.011 |
| PerceiverCPI | 0.848±0.011 | 0.826±0.012 | 0.724±0.010 | 0.714±0.011 | 0.708±0.012 | 0.728±0.010 | 0.268±0.010 |
| DrugBAN | 0.852±0.010 | 0.830±0.011 | 0.718±0.011 | 0.708±0.012 | 0.702±0.013 | 0.720±0.011 | 0.260±0.011 |
| CAT-DTI | 0.895±0.008 | 0.889±0.009 | 0.784±0.009 | 0.774±0.010 | 0.778±0.010 | 0.758±0.009 | 0.286±0.009 |
| GraphBAN | 0.920±0.005 | <u>0.904±0.006</u> | 0.790±0.007 | <u>0.782±0.008</u> | 0.772±0.008 | <u>0.798±0.007</u> | 0.313±0.007 |
| ExplainBind | **0.940±0.004** | **0.926±0.005** | **0.805±0.006** | **0.793±0.007** | **0.788±0.007** | **0.810±0.006** | **0.330±0.007** |
| **InteractBind-P (31% OOD)** |  |  |  |  |  |  |  |
| MolTrans | 0.785±0.015 | 0.755±0.017 | 0.670±0.012 | 0.660±0.013 | 0.653±0.015 | 0.675±0.011 | 0.228±0.013 |
| TransformerCPI | 0.806±0.014 | 0.774±0.015 | 0.684±0.012 | 0.674±0.013 | 0.668±0.014 | 0.688±0.012 | 0.236±0.012 |
| HyperAttentionDTI | 0.805±0.013 | 0.789±0.014 | 0.696±0.011 | 0.686±0.012 | 0.680±0.013 | 0.700±0.011 | 0.246±0.011 |
| PerceiverCPI | 0.832±0.011 | 0.816±0.012 | 0.716±0.010 | 0.706±0.011 | 0.700±0.012 | 0.720±0.010 | 0.262±0.010 |
| DrugBAN | 0.842±0.010 | 0.820±0.012 | 0.710±0.011 | 0.700±0.012 | 0.695±0.013 | 0.712±0.011 | 0.255±0.011 |
| CAT-DTI | 0.879±0.008 | 0.874±0.009 | 0.744±0.009 | 0.734±0.010 | 0.728±0.010 | 0.748±0.009 | 0.279±0.009 |
| GraphBAN | <u>0.910±0.006</u> | <u>0.894±0.007</u> | <u>0.780±0.008</u> | <u>0.768±0.009</u> | <u>0.764±0.009</u> | <u>0.784±0.008</u> | <u>0.306±0.008</u> |
| ExplainBind | **0.930±0.005** | **0.915±0.006** | **0.795±0.007** | **0.783±0.007** | **0.778±0.008** | **0.800±0.007** | **0.322±0.007** |
| **InteractBind-P (28% OOD)** |  |  |  |  |  |  |  |
| MolTrans | 0.768±0.016 | 0.740±0.017 | 0.658±0.013 | 0.648±0.014 | 0.640±0.016 | 0.660±0.012 | 0.225±0.013 |
| TransformerCPI | 0.781±0.014 | 0.756±0.015 | 0.672±0.012 | 0.662±0.013 | 0.655±0.014 | 0.676±0.012 | 0.229±0.012 |
| HyperAttentionDTI | 0.788±0.013 | 0.772±0.014 | 0.684±0.011 | 0.674±0.012 | 0.668±0.013 | 0.688±0.011 | 0.239±0.011 |
| PerceiverCPI | 0.814±0.011 | 0.801±0.012 | 0.706±0.010 | 0.696±0.011 | 0.690±0.012 | 0.710±0.010 | 0.255±0.010 |
| DrugBAN | 0.825±0.010 | 0.805±0.011 | 0.700±0.011 | 0.688±0.012 | 0.682±0.013 | 0.705±0.011 | 0.250±0.011 |
| CAT-DTI | 0.859±0.008 | 0.868±0.009 | 0.732±0.009 | 0.722±0.010 | 0.716±0.010 | 0.736±0.009 | 0.272±0.009 |
| GraphBAN | <u>0.900±0.006</u> | <u>0.882±0.007</u> | 0.768±0.008 | <u>0.758±0.009</u> | 0.750±0.009 | <u>0.774±0.008</u> | 0.296±0.008 |
| ExplainBind | **0.910±0.005** | **0.895±0.006** | **0.782±0.007** | **0.770±0.007** | **0.765±0.008** | **0.788±0.007** | **0.310±0.007** |
| **InteractBind-P (25% OOD)** |  |  |  |  |  |  |  |
| MolTrans | 0.715±0.020 | 0.675±0.022 | 0.622±0.015 | 0.612±0.016 | 0.608±0.018 | 0.624±0.014 | 0.210±0.015 |
| TransformerCPI | 0.718±0.017 | 0.696±0.019 | 0.634±0.014 | 0.624±0.015 | 0.618±0.016 | 0.638±0.014 | 0.220±0.014 |
| HyperAttentionDTI | 0.752±0.016 | 0.710±0.018 | 0.646±0.013 | 0.636±0.014 | 0.630±0.015 | 0.650±0.013 | 0.228±0.014 |
| PerceiverCPI | 0.779±0.014 | 0.738±0.016 | 0.668±0.012 | 0.658±0.013 | 0.652±0.014 | 0.672±0.012 | 0.242±0.013 |
| DrugBAN | 0.781±0.011 | 0.775±0.014 | 0.689±0.013 | 0.678±0.012 | 0.670±0.015 | 0.715±0.012 | 0.235±0.013 |
| CAT-DTI | 0.801±0.011 | 0.800±0.013 | 0.714±0.011 | 0.704±0.012 | 0.708±0.013 | 0.706±0.011 | 0.252±0.012 |
| GraphBAN | <u>0.832±0.010</u> | <u>0.803±0.003</u> | <u>0.742±0.003</u> | <u>0.738±0.013</u> | <u>0.722±0.014</u> | <u>0.754±0.012</u> | <u>0.265±0.012</u> |
| ExplainBind | **0.850±0.006** | **0.836±0.008** | **0.765±0.008** | **0.754±0.009** | **0.748±0.010** | **0.770±0.009** | **0.280±0.009** |

## Footnotes

1 https://pytorch.org/

